# Hierarchical TAF1-dependent co-translational assembly of the basal transcription factor TFIID

**DOI:** 10.1101/2023.04.05.535704

**Authors:** Andrea Bernardini, Pooja Mukherjee, Elisabeth Scheer, Ivanka Kamenova, Simona Antonova, Paulina Karen Mendoza Sanchez, Gizem Yayli, Bastien Morlet, H.T. Marc Timmers, László Tora

## Abstract

Large heteromeric multiprotein complexes play pivotal roles at every step of gene expression in eukaryotic cells. Among them, the 20-subunit basal transcription factor TFIID nucleates RNA polymerase II preinitiation complex at gene promoters. Here, by combining systematic RNA-immunoprecipitation (RIP) experiments, single-molecule imaging, proteomics and structure-function analyses, we show that TFIID biogenesis occurs co-translationally. We discovered that all protein heterodimerization steps happen during protein synthesis. We identify TAF1 – the largest protein in the complex – as a critical factor for TFIID assembly. TAF1 acts as a flexible scaffold that drives the co-translational recruitment of TFIID submodules preassembled in the cytoplasm. Altogether, our data suggest a multistep hierarchical model for TFIID biogenesis that culminates with the co-translational assembly of the complex onto the nascent TAF1 polypeptide. We envision that this assembly strategy could be shared with other large heteromeric protein complexes.

## INTRODUCTION

A considerable portion of genes produce proteins that become part of multiprotein complexes. The function and structure of many of these molecular machineries are being extensively investigated, however the understanding of the precise steps guiding their assembly process is of key importance to interpret physio-pathological perturbations of their components and grasp how cells work.

Large heteromeric protein complexes are implicated in all aspects of gene expression and uncovering their assembly mechanism is particularly challenging. Several different subunits – synthesized by separate mRNA molecules – must productively interact with their direct partner/s in the crowded cellular environment and sequentially build larger assemblies, while minimizing off-pathway interactions and aggregation.

A fascinating mechanism thought to facilitate the formation of protein complexes is co-translational assembly (here abbreviated as Co-TA), whereby the newly synthesized nascent protein chain establishes the interaction with the partner before being released from the ribosome^1, 2^. Co-TA can be sequential (also termed directional), when a fully translated protein is recruited to the partner nascent chain, or simultaneous (also termed symmetrical), if both interactors are nascent chains. Coupling specific subunit-subunit assembly with translation would reduce the exposure of aggregation-prone domains, facilitate the formation of intricate protein-protein interfaces and impart a sequential order for the assembly of different subunits^1, 3^. Co-TA has been demonstrated to partake in the heterodimerization of several yeast proteins^4–7^, including the localized assembly of specific subunits of the massive nuclear pore complex^8, 9^.

A key regulatory step in gene expression is transcription initiation. Including RNA polymerases, most of the molecular machineries that act on transcription initiation are large heteromeric protein complexes. Among them, the ∼1.3 MDa basal transcription factor TFIID makes contacts with core promoter DNA elements, promotes TATA-binding protein (TBP) loading on upstream DNA and works as a dynamic scaffold for the formation of RNA polymerase II preinitiation complex (PIC) on all protein-coding genes^10–12^.

In metazoans, TFIID is constituted by TBP and 13 different TBP-associated factors (TAFs)^13^. Since the first attempts to define TFIID structure, it was clear that the complex could be subdivided in three lobes (**Fig. 1a**)^14, 15^. More recently, single-particle cryo-electron microscopy (cryo-EM) models of yeast and human TFIID shed light on the position and atomic interactions among its subunits^10, 16–18^.

**Figure 1.**
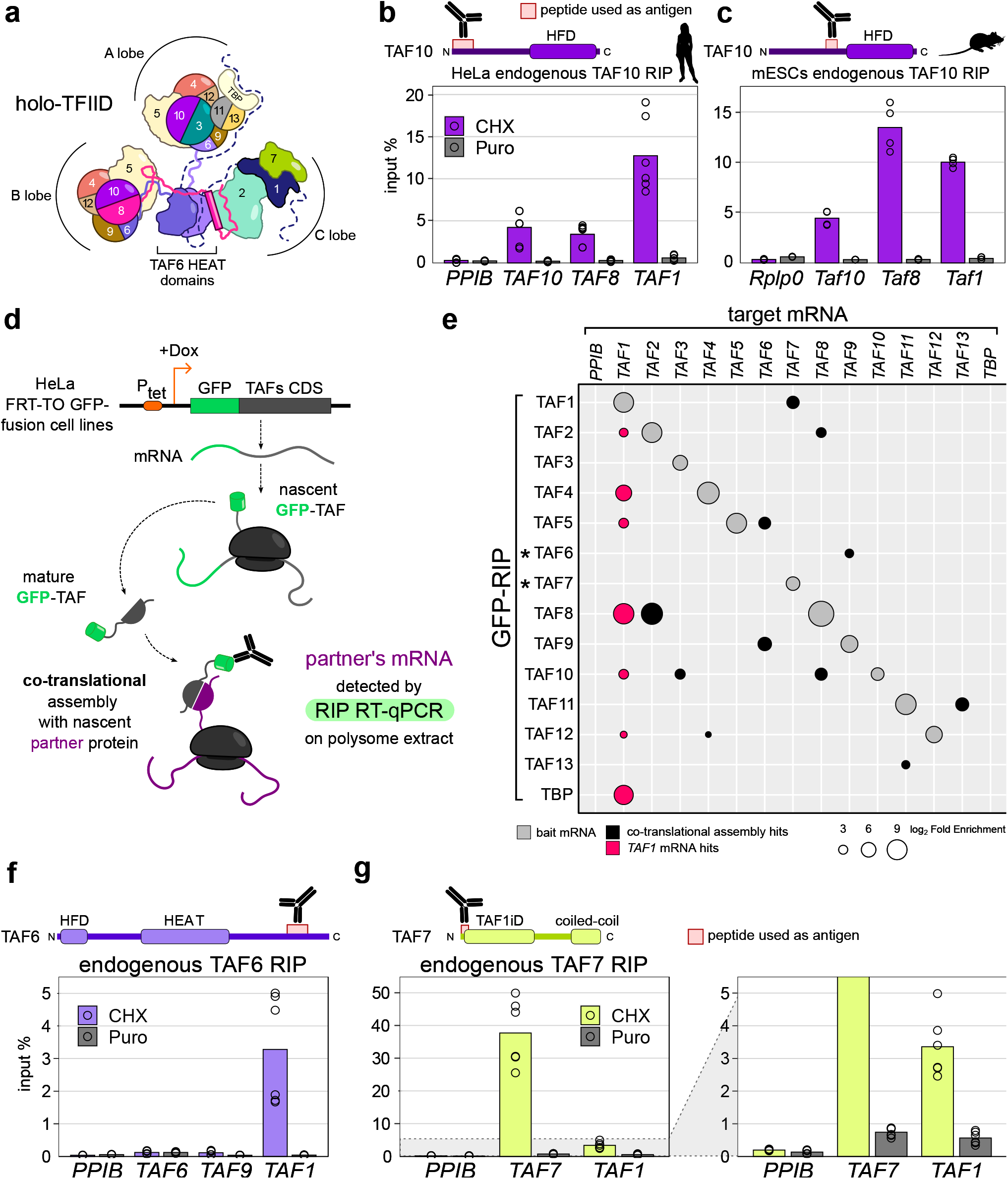
A systematic assay expands the network of co-translational interactions in TFIID and identifies nascent TAF1 polypeptide as a central hub in the assembly process. **a**, Schematic structure of the three lobed holo-TFIID complex. Half-circle subunits correspond to HFD-containing partners. TAF1 is represented with a dotted line. **b**, RNA immunoprecipitation (RIP) assays using an antibody against endogenous human TAF10 performed on HeLa cell polysome extracts. Potential target mRNAs were tested by RT-qPCR. Data points correspond to technical duplicates from three biological replicates. **c**, RIP coupled RT-qPCR assays using an antibody against endogenous mouse TAF10 performed on mESCs. Data points correspond to technical duplicates from two biological replicates. **d**, Schematic representation of the GFP-RIP coupled RT-qPCR assay using different HeLa cell lines expressing doxycycline (Dox) inducible GFP-TAFs to systematically probe co-translational assembly events between TFIID subunits. **e**, Matrix summarizing the results of the systematic GFP-RIP coupled RT-qPCR assay schematized in panel (d). Each GFP-tagged TFIID subunit was used as bait in a GFP-RIP assay from polysome extracts (rows) and enrichment for TFIID subunits mRNAs was assessed by RT-qPCR (columns). Circles area is proportional to mRNA log2 fold enrichment (FE) over mock IP. Combinations with less than 4-fold the FE of negative control target mRNA (*PPIB*) are not shown in the plot and considered negative. Grey circles represent hits for baits’ mRNA. Black circles represent co-translational assembly hits. Red circles highlight the widespread enrichment for *TAF1* mRNA from RIP of several TFIID subunits. Stars refer to subunits for which GFP fusion resulted in ambiguous protein functionality. Results represent the mean of two biological replicates. **f**, RIP coupled RT-qPCR assays using an antibody against endogenous TAF6 performed on HeLa cells. Note that the C-terminal location of the region recognized by the TAF6 antibody prevented the detection of the nascent TAF6 protein along with its own mRNA and the simultaneous Co-TA with TAF9 (the HFD partner of TAF6). **g**, RIP coupled RT-qPCR assays using an antibody against endogenous TAF7 performed on HeLa cells. Antigen region for antibodies used in the assay are indicated. Data points correspond to technical triplicates from two biological replicates. HFD: histone-fold domain; HEAT: HEAT-repeat domain; TAF1iD: TAF1-interaction domain; Cycloheximide (CHX) “freezes” translating ribosomes, with the nascent polypeptides, on the mRNA. In contrast, puromycin (Puro) induces premature nascent polypeptide chain termination and release from ribosome/mRNA.

Nine TAFs contain a histone-fold domain (HFD) that dictates five defined dimerization interfaces within the complex, namely TAF4/TAF12, TAF6/TAF9, TAF3/TAF10, TAF8/TAF10, TAF11/TAF13. A set of five TAFs (TAF4/TAF5/TAF6/TAF9/TAF12) – named core-TFIID – are present in two copies, constituting a pseudo-symmetrical unit within the complex that occupies both A and B lobes. These two lobes differ in the dimerization partner of TAF10: TAF3 in A lobe and TAF8 in B lobe, respectively. Moreover, lobe A is characterized by the additional TAF11/TAF13 HFD pair, which bridges TBP to the rest of the lobe (**Fig. 1a**). Finally, the C lobe is constituted by the structured domains of the TAF1/TAF7 dimer, TAF2 and the central HEAT domains of the two copies of TAF6. B and C lobes are connected through the conserved C-terminal portion of TAF8 which directly interacts with TAF2^19^, making the composite B/C lobe rigid, while A lobe remains flexibly connected to the rest of the complex through TAF6 linker region.

How and where TFIID assembles in cells is a longstanding question which has been surprisingly overlooked. Classically, the holo-complex is isolated from nuclear extracts, while attempts to isolate endogenous assemblies in the cytoplasm led to the identification of preformed TAF2/TAF8/TAF10 and TAF11/TAF13 modules^20, 21^. Another hint on the formation of cytoplasmic TFIID submodules came from the isolation of a stable TAF5/TAF6/TAF9 complex using an inducible cellular expression system^22^. These observations led to a model whereby different TFIID modules are formed in the cytoplasm and independently translocate in the nucleus, where holo-complex formation would take place^23^.

Supporting the early steps of TFIID biogenesis in the cytoplasm, we demonstrated Co-TA events between three pairs of TFIID subunits, either sequentially (TAF10>TAF8, TBP>TAF1) or simultaneously (TAF6/TAF9)^24^. With the aim of searching for additional Co-TA events in TFIID, we carried out a broad combination of complementary approaches, including systematic RIPs, imaging and proteomics. During the course of our analyses, we identified a series of novel pairwise Co-TA events that shape the early steps of TFIID assembly. Unexpectedly, we uncovered a new role for TAF1 nascent protein as a co-translational ‘landing platform’ for other preassembled TFIID submodules. Thus, our data unravel the assembly pathway of the entire TFIID complex, whose principles might be applicable to the biogenesis of other large multiprotein complexes.

## RESULTS

### A systematic RIP assay expands the network of co-translational interactions in TFIID and identifies nascent TAF1 polypeptide as a central hub in the assembly process

When reanalyzing our previously published TAF10 RNA immunoprecipitation (RIP)-microarray data^24^ by using recent genomic annotations, *TAF1* mRNA scored as a positive hit (**Extended Data Fig. 1a**). Intriguingly, TAF1 is devoid of HFDs and it is not known to directly interact with TAF10 within TFIID, raising the possibility that higher-order co-translational interactions might take place. This surprising observation prompted us to perform TAF10 RIP-qPCR on HeLa cells polysome extracts (**Fig. 1b**). RIPs are performed in the presence of cycloheximide, which “freezes” translating ribosomes with the nascent polypeptides on the mRNA to prevent their dissociation throughout the protocol. In contrast, puromycin was used to induce premature nascent chain termination, to exclude direct protein/RNA interactions. Indeed, we detected a strong enrichment of *TAF1* mRNA in TAF10 RIPs, along with the expected mRNAs of *TAF10* itself and its HFD partner *TAF8*. *TAF1* mRNA enrichment was reproducible and puromycin-sensitive, suggesting a co-translational association of TAF10 with the nascent TAF1 polypeptide. To rule out biases from the cellular system or the antibody used, we performed the same experiment on E14 mouse embryonic stem cells (mESCs) using a different monoclonal antibody than the one used in the human system. We found that the anti-TAF10 RIP enriched *Taf1* mRNA also in this mouse cellular system (**Fig. 1c**), pointing at a conserved phenomenon.

*TAF1* mRNA enrichment in TAF10 RIPs was unexpected, thus we wanted to assess whether this observation was TAF10-specific or more global in TFIID. To systematically assess all co-translational assembly events within TFIID complex, we used a series of inducible HeLa cell lines engineered to express each TFIID subunit as a fusion protein with a N-terminal GFP tag^25^. These GFP-tagged TAFs were shown to incorporate in TFIID purified from nuclear extracts^26^. We performed GFP-RIP assays on polysome extracts for each individual TFIID subunit and systematically tested for enrichment of mRNAs encoding all the TFIID subunits by RT-qPCR (**Fig. 1d**). The results of this systematic RIP-qPCR screening (**Extended Data Fig. 1b**) are summarized in **Fig. 1e**. These assays confirmed the previously published TFIID Co-TA subunit pairs (TAF10>TAF8, TAF6/TAF9 and TBP>TAF1), thereby validating the general reliability of the system (**Fig. 1e**). Strikingly, our systematic assay revealed that *TAF1* mRNA was enriched in IPs of several distinct TFIID subunits (red circles in **Fig. 1e**). In this systematic analysis, TAF10 RIP also scored positive for *TAF1*, confirming the observations from endogenous TAF10 RIPs (**Fig. 1b-c**). Apart from TAF10, also TAF2, TAF4, TAF5, TAF8, TAF12 and TBP RIPs retrieved *TAF1* mRNA (**Fig. 1e**).

In addition, novel subunit pairs undergoing Co-TA were detected (**Table S1**). Most of them include well-established HFD partners: TAF10 interacts co-translationally also with nascent TAF3; TAF12 with nascent TAF4; and TAF11/TAF13 are reciprocally enriched, hinting at symmetrical Co-TA. Our systematic RIP assay also revealed Co-TA among direct partner subunits that do not interact through a HFD. For instance, TAF2 and TAF8, which are known to directly interact in TFIID (**Fig. 1a** and **Extended Data Fig. 1c**), reciprocally enriched the partner’s mRNA suggesting simultaneous Co-TA. TAF5 enriched *TAF6* mRNA, one of its most intimate interactors within core-TFIID, where TAF6 contributes with a β-strand to the last blade of TAF5 WD40 β-propeller domain (**Extended Data Fig. 1d**). Finally, TAF1 enriched the mRNA of *TAF7*, its most well characterized direct partner within TFIID (**Fig. 1e**).

We noted for three of the GFP-fusion protein expressing cell lines that we could not retrieve the bait mRNA in our RIP assays (TAF6, TAF13 and TBP) and that the anti-GFP-TAF7 RIP failed to retrieve *TAF1*. Note that TBP/TAF1 Co-TA was already shown with endogenous TBP RIPs in our previous report^24^. To complete our systematic screening, we performed RIPs with antibodies recognizing endogenous TAF6 and TAF7 and observed a robust puromycin-sensitive enrichment of *TAF1* mRNA in both TAF6 and TAF7 RIPs (**Fig. 1f-g** and **Extended Data Fig. 1e**). Overall, these observations expand the repertoire of TFIID subunits that follow a co-translational pathway for the assembly with their partners, and importantly identify the nascent TAF1 protein as a potential hub for the recruitment and assembly of many TFIID subunits.

### Endogenous TFIID subunits are localized in physical proximity to *TAF1* mRNA in the cytoplasm of human cells

Our RIP-based observations suggest that during *TAF1* mRNA translation, several TFIID subunits physically associate with TAF1 nascent polypeptide. To physically localize and quantify these events in an endogenous cellular context, we combined immunofluorescence (IF) against several TFIID subunits with single-molecule RNA fluorescence *in situ* hybridization (smFISH) using HeLa cells.

First, we applied this strategy to detect TAF1 nascent protein and estimate the fraction of actively translated *TAF1* mRNAs. To this end we used an IF-validated TAF1 antibody recognizing a N-terminal antigen and combined it with *TAF1* mRNA smFISH (**Fig. 2a**). We used *CTNNB1* as a negative control target mRNA in smFISH. The average number of cytoplasmic mRNAs per cell for *TAF1* was ∼16 and about 120 for *CTNNB1* (**Fig. 2b**). Next, we combined TAF1 IF with *TAF1* or *CTNNB1* smFISH (**Fig. 2c**) and quantified the number of *TAF1* mRNA molecules co-localizing with TAF1 protein spots in confocal microscopy images. On average, ∼55% of *TAF1* cytoplasmic mRNAs co-localized with TAF1 IF spots (**Fig. 2d**). The co-localized fraction decreased more than ten-fold upon puromycin treatment, proving a dependence on mRNA/ribosome/nascent chain integrity. The very low fraction (∼1%) of co-localization with *CTNNB1* mRNA was puromycin-insensitive and represents a baseline of random co-localization. We also validated the specificity of the *TAF1* smFISH signal by siRNA-mediated knockdown of *TAF1* (**Extended Data Fig. 2**). Hence, this experimental strategy is able to detect co-translational events and we could estimate that roughly half of *TAF1* mRNAs are detected as being actively translated in HeLa cells. Note however, that transcripts that are not detected co-localizing with protein spots could also be translated, yet to an extent that does not allow detection of the corresponding protein, or protein partners.

**Figure 2.**
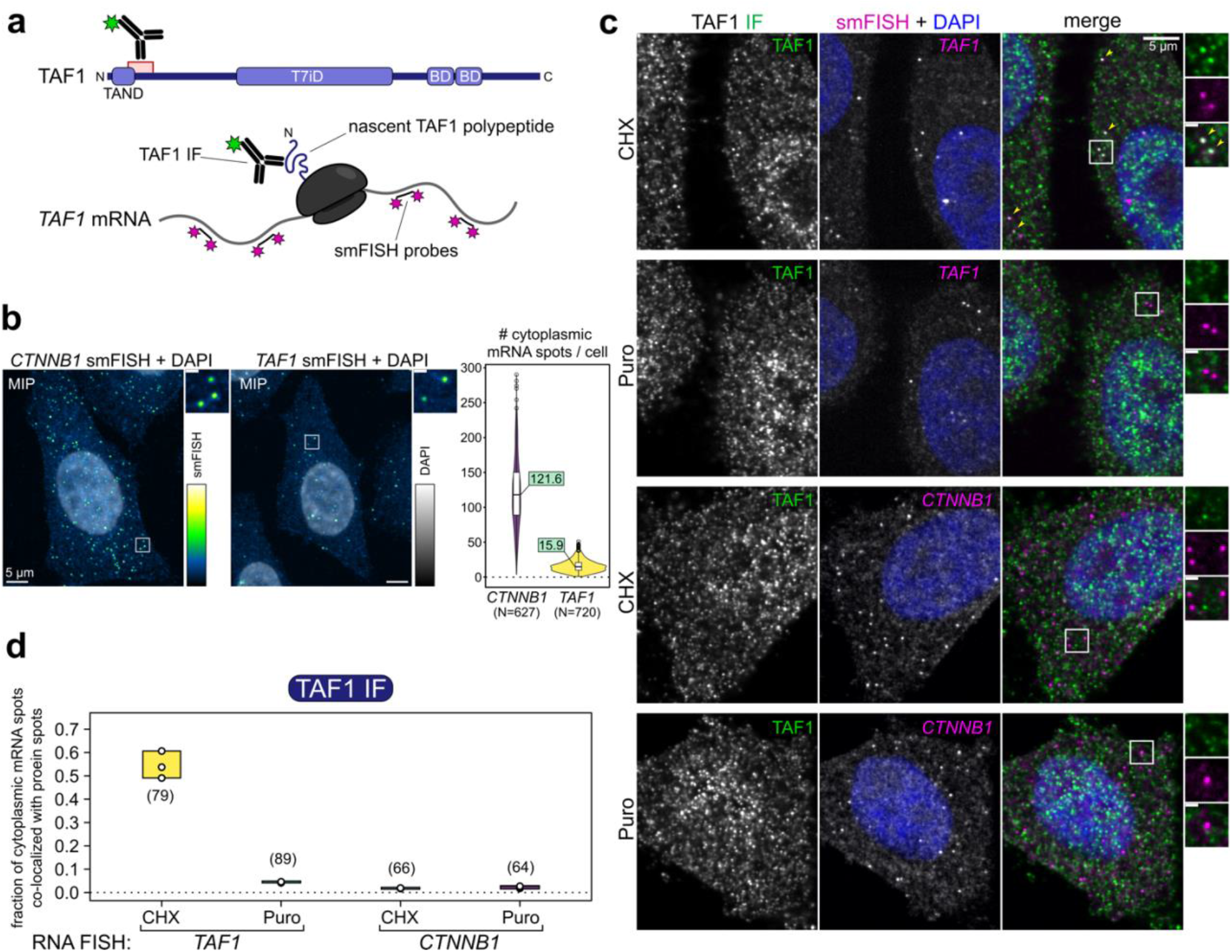
Endogenous nascent TAF1 protein detection. **a**, Schematic overview of the imaging strategy to detect actively translated *TAF1* endogenous mRNAs with a combination of single-molecule RNA FISH (smFISH) and immunofluorescence (IF). The antigen region recognized by the TAF1 antibody used in the assay is indicated. TAND: TAF1 N-terminal domain; T7iD: TAF7-interaction domain; BD: bromodomain. **b**, Representative confocal maximum intensity projections (MIPs) of smFISH against the negative control *CTNNB1* and *TAF1* mRNAs in HeLa cells. The plot on the right shows the absolute number of cytoplasmic mRNAs per cell (mean values are reported in the boxes, total number of cells in brackets). **c**, Representative multicolor confocal images for the co-localization assay shown in (a). Each image is a single confocal optical slice. TAF1 protein IF and *TAF1* mRNA detection in the merged image are shown in green and magenta, respectively. Colocalizing spots are indicated with yellow arrows. Zoom-in regions (white squares) are shown on the right. CHX: cycloheximide; Puro: puromycin (see also the legend of Fig. 1). Scale bars in the insets corresponds to 1 μm. **d**, Quantification of the fraction of mRNAs co-localized with protein signal for each experimental condition. Each open circle corresponds to an independent field of view (N = 3, total number of cells is in brackets).

Next, we applied the same strategy to subunits belonging to distinct TFIID lobes to assess their spatial proximity with *TAF1* mRNA, as would be predicted if they associate with nascent TAF1 polypeptide during translation (**Fig. 3a**). First, we assessed the combination with TBP (lobe A component) (**Fig. 3b**), as its co-translational association with TAF1 was already dissected^24^. We found that ∼6% of cytoplasmic *TAF1* mRNAs co-localized with TBP, while less than 1% of the negative control *CTNNB1* mRNAs did so. Moreover, the fraction of co-localized TAF1 mRNAs robustly reduced upon puromycin treatment, confirming the Co-TA between the two subunits.

**Figure 3.**
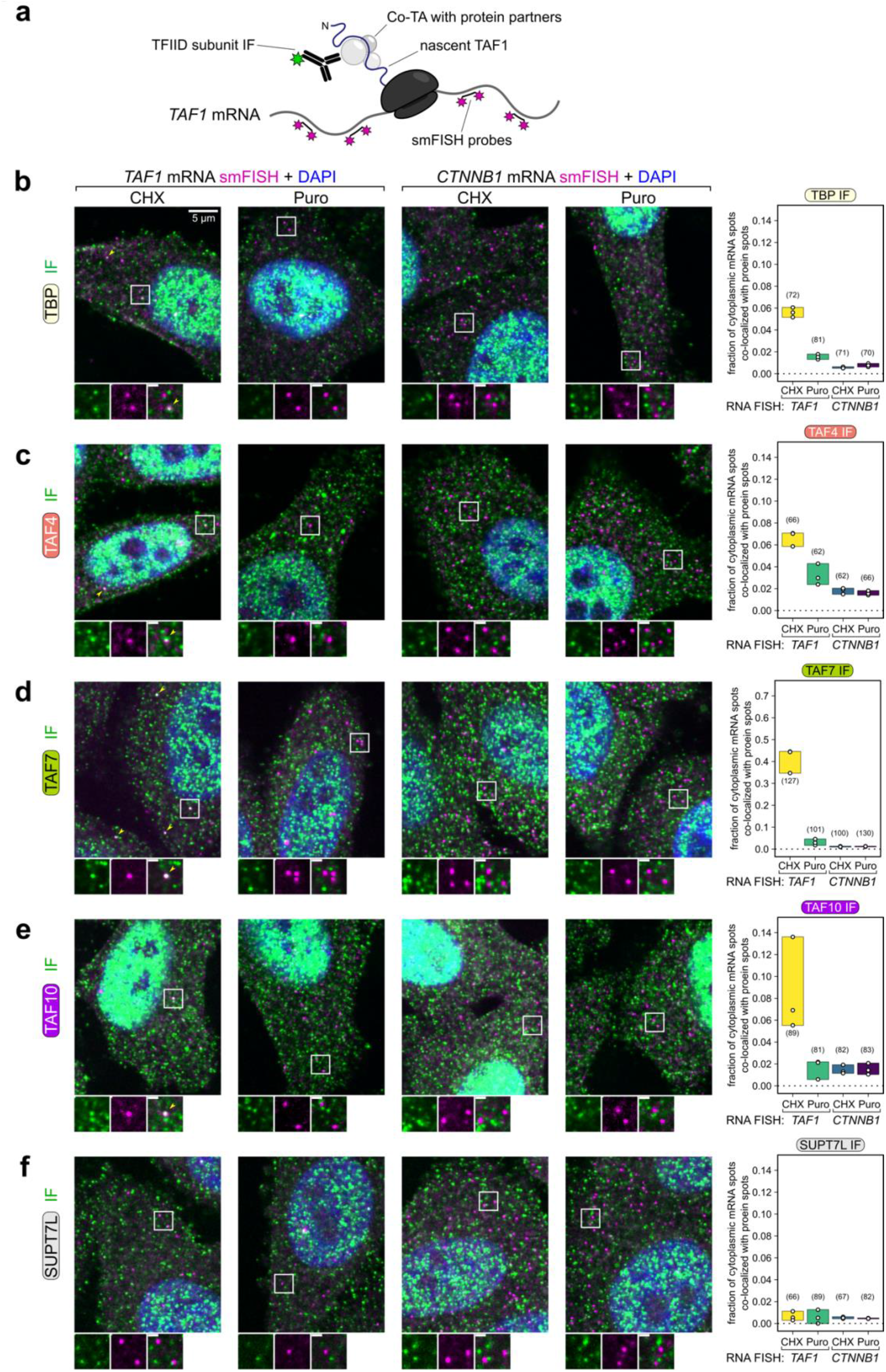
Endogenous TFIID subunits are localized in physical proximity to TAF1 mRNA in the cytoplasm of human cells. **a**, Schematic overview of the imaging strategy to detect Co-TA events of endogenous TFIID subunits on *TAF1* mRNA with a combination of single-molecule RNA FISH (smFISH) and immunofluorescence (IF). **b-f**, Representative multicolor confocal images for the co-localization assay shown in a for each assessed subunit. Each image is a single multichannel confocal optical slice. CHX: cycloheximide; Puro: puromycin. Protein IF and *TAF1* mRNA detection are shown in green and magenta, respectively. Colocalizing spots are indicated with yellow arrows. Zoom-in regions (white squares) are shown below each image. The scale bar in the insets corresponds to 1 μm. The plots on the right report the fraction of target mRNAs co-localized with protein signal for each experimental condition. Each open circle corresponds to an independent field of view (N = 3, total number of cells is in brackets).

We then performed the experiment with IFs against TAF4 (part of core-TFIID, component of lobes A and B), TAF7 (lobe C component) and TAF10 (lobes A and B component). All tested TAFs positively co-localized with *TAF1* mRNA (**Fig. 3c-e**). TAF4 (**Fig. 3c**) and TAF10 (**Fig. 3e**) co-localization levels with *TAF1* were comparable with those of TBP (**Fig. 3b**), while for TAF7 (**Fig. 3d**) they were considerably higher, with ∼40% of *TAF1* mRNAs co-localizing with TAF7 spots. Although we noticed a partial puromycin resistance for TAF4, puromycin treatment consistently reduced the fraction of co-localization for all TAFs. Instead, *TAF1* co-localization with SUPT7L – an unrelated HFD subunit of the SAGA complex – was very low (<1%) and not affected by puromycin (**Fig. 3f**), thus confirming the specificity of the results. We then simultaneously probed cells for TBP and TAF7 subunits by dual-color IF (**Extended Data Fig. 3a**). We found that ∼50% of

TBP-positive *TAF1* RNA spots were simultaneously co-localized with TAF7. Puromycin treatment drastically reduced the frequency of co-localization, nearly abolishing the double-positive events (**Extended Data Fig. 3b**). Overall, these microscopy observations support our systematic RIP-qPCR screening data, further suggesting that multiple TFIID subunits are recruited on TAF1 nascent polypeptide during the protein synthesis of the latter.

### The cytoplasm is populated by multisubunit TFIID ‘building blocks’

The novel Co-TA events revealed by our RIP-screening mapped to well characterized, direct partners within TFIID, such as TAF2/TAF8, TAF3/TAF10, TAF1/TAF7 and others^10, 18^, substantiating our observations. Surprisingly, this is not the case for the majority of the nine Co-TA events directed on nascent TAF1 polypeptide (**Fig. 1e-g**). Specifically, no direct TAF1 interactions are known with TAF4, TAF5, TAF8, TAF10 and TAF12, although novel interaction domains cannot be excluded, since only a subset of TAF1 residues has been mapped in the published cryo-EM maps^10, 18^. To better understand how the above discovered Co-TA events may participate in TFIID assembly, we set out to analyze the composition of potential TFIID assemblies in the cytoplasm of human cells. To this end, we immunopurified endogenous TFIID subunits from HeLa cytoplasmic extracts and analyzed the immunoprecipitated endogenous complexes by label-free mass-spectrometry (**Fig. 4a-f**). The same immunoprecipitations (IPs) invariably retrieved holo-TFIID from nuclear extracts, confirming the effectiveness of the antibodies used (**Extended Data Fig. 4a**).

**Figure 4.**
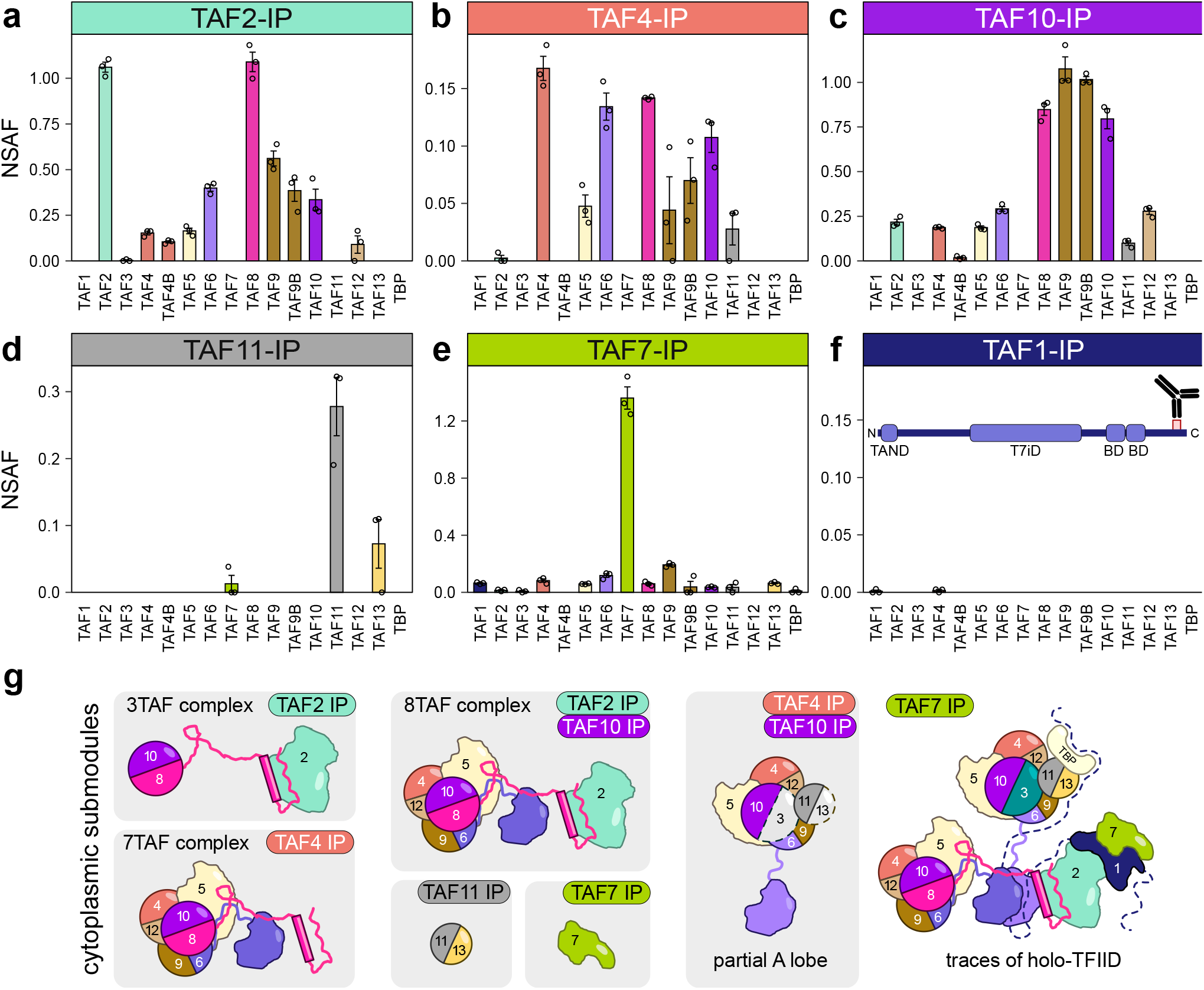
The cytoplasm is populated by multisubunit TFIID ‘building blocks’. **a-f**, Immunoprecipitation (IP) of endogenous TFIID subunits coupled to label-free mass-spectrometry performed on human HeLa cells cytoplasmic extracts. Bar plots represent the average NSAF (normalized spectral abundance factor) value for each detected subunit in technical triplicates. The antigen position of TAF1 antibody used for IP is shown in panel (f). **g**, Visual summary of cytoplasmic TFIID submodules inferred from IP-MS data. IPs that enriched the given submodule are indicated.

The immunopurified cytoplasmic TAF2 was abundantly associated with TAF8, in accordance with their Co-TA (**Fig. 1e**), and partially integrated in the so called 8TAF complex (composed of the core-TFIID + TAF2/TAF8/TAF10, see Introduction; **Fig. 4**)^21, 27^. The majority of immunopurified cytoplasmic TAF4 was complexed with TAF8, TAF6, TAF9/9B, TAF10 and TAF5 (ordered by NSAF values, **Fig. 4b**), that we interpreted as the 7TAF complex (core-TFIID + TAF8/TAF10, note that the missing detection of TAF12 could be due to the documented post-translational modifications of this subunit). The presence of low amounts of TAF11 copurified with TAF4 hinted at the incorporation of the latter in what could be a partial A lobe. Interestingly, in the cytoplasmic anti-TAF4 IPs we find only TAF4, while in the nuclear anti-TAF4 IP we find peptides from TAF4 and its paralog TAF4B (**Fig. 4b** and **Extended Data Fig. 4a**). These findings suggest that the isolated cytoplasmic TAF4-containing building block is either lobe A, or lobe B (as indicated in **Fig. 4g**), containing only one copy of TAF4. In contrast, the detection of both TAF4 and TAF4B in nuclear TAF4 IP suggests the isolation of the holo-TFIID, where lobes A and B are part of the same complex.

Endogenous cytoplasmic TAF10 IP retrieved similar amounts of the HFD-partner TAF8, with a relevant portion of the heterodimer associated with core-TFIID and TAF2 in the 8TAF complex (**Fig. 4c**). In this case, the low amounts of TAF11 also hint at a fraction of TAF10 incorporated in a partially assembled A lobe. On the other hand, we found TAF11 partially associated with its natural partner TAF13 in the TAF11 IP^20^, with no detectable amounts of other A lobe subunits (**Fig. 4d**). Most immunopurified TAF7, the direct TAF1 partner in TFIID, was non-complexed (**Fig. 4e**). Note however that cytoplasmic TAF7 co-purified with trace amounts of TFIID subunits – including TAF1 (**Extended Data Fig. 4b**). Interestingly, in contrast to nuclear extracts (**Extended Data Fig. 4a**), cytoplasmic TAF1 IP did not enrich any TFIID component, with the bait itself barely detectable (**Fig. 4f**). Given the IP-grade TAF1 antibody recognizing a C-terminal epitope along the protein, we conclude that the abundance of TAF1 mature protein in the cytoplasm is below the detection limit in this analysis.

Together, these results demonstrate that the cytoplasm of HeLa cells is populated by different multisubunit TFIID submodules, likely representing stable intermediates along the assembly pathway of the complex (**Fig. 4g**). None of the cytoplasmic IPs, except for TAF7, co-purified TAF1, suggesting that TAF1 is present in low amounts in the cytoplasm and is the limiting factor in TFIID assembly. These findings further point to a co-translational recruitment mechanism where the pre-assembled TFIID ‘building blocks’ would associate with nascent TAF1 polypeptide, in good agreement with our RIP-RT-qPCR and imaging experiments.

### Three crosslinking hotspots identified on TAF1 correspond to distinct anchor points for specific TFIID building blocks

TAF1 is the biggest subunit of TFIID (1872 aa) and only ∼47% of the protein structure has currently been solved by experimental means. To rationalize how nascent TAF1 polypeptide could work as a hub for TFIID assembly, we analyzed all available crosslinking mass spectrometry (X-linking MS) experiments performed on highly purified TFIID or PIC-incorporated TFIID, including ours^10, 18, 19^ (**Table S2**). The intercrosslinks between TAF1 and other TFIID subunits detected in at least two independent datasets are summarized in **Fig. 5a**. From the N- to C-terminal regions, we could isolate three main proximity/crosslinking ‘hotspots’ along TAF1: (1) a loose region crosslinked with TBP and its interacting partners TAF11/TAF13, (2) a well-defined hotspot rich in crosslinks with TAF6 along with single positions recurrently associated with TAF5, TAF8 and TAF9, and (3) a large central region found extensively cross-linked to TAF7 and – to a lesser extent – to TAF2. The combination of the crosslinking hotspots (**Fig. 5a**) with TAF1 sequence features (conservation, structural disorder, **Fig. 5b**), annotated functional domains (**Fig. 5c**) and structural observations (**Fig. 5d**) shows that TAF1 is a flexible scaffold protein that connects all TFIID submodules by three main anchor points (described below).

**Figure 5.**
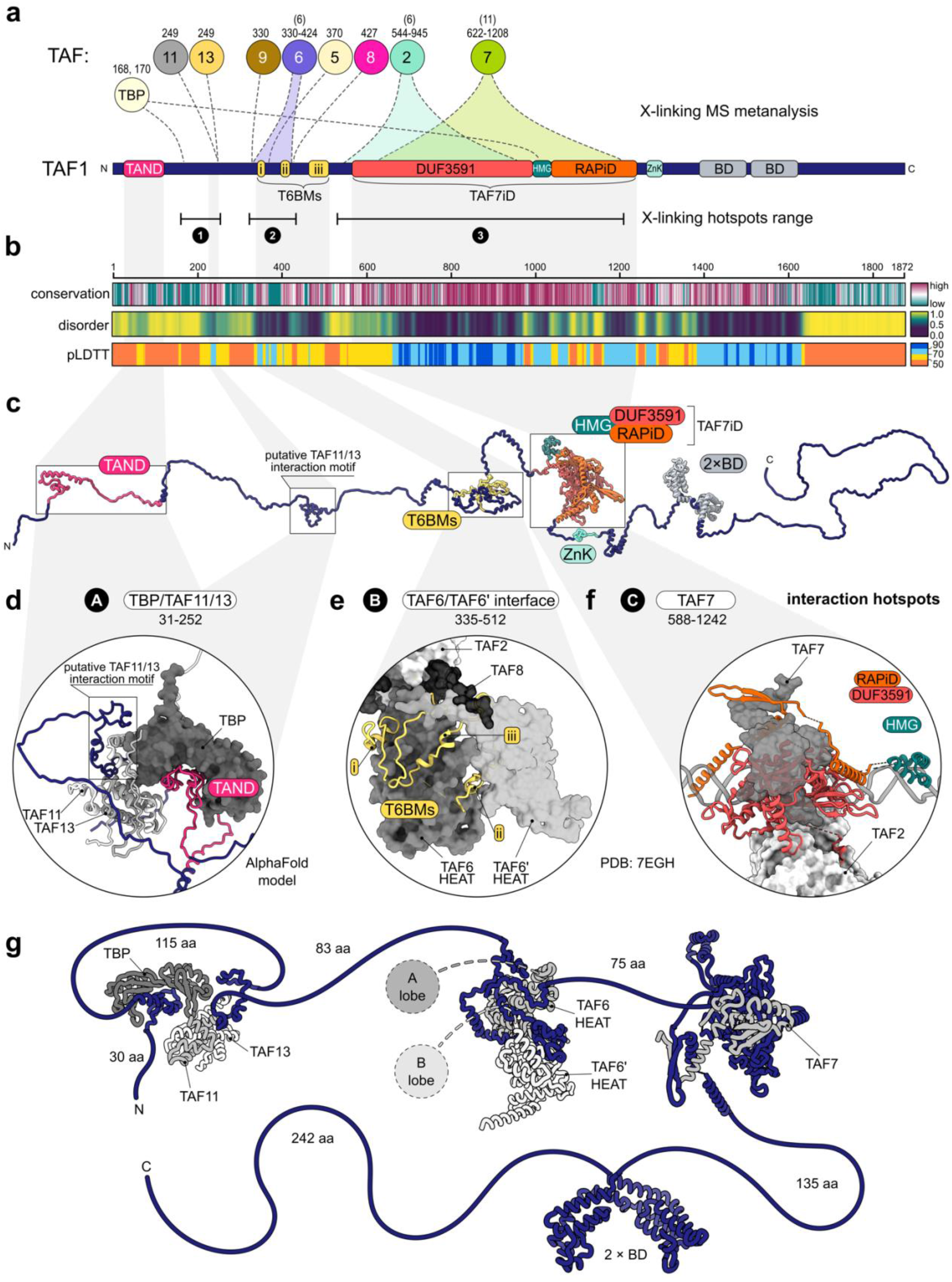
Three crosslinking hotspots identified on TAF1 correspond to distinct anchor points for specific TFIID building blocks. **a**, Summary of TAF1-centred crosslinking-mass spectrometry metanalysis derived from three independent studies^10, 18, 19^. Inter-protein crosslinks between TAF1 and other TFIID subunits (TAFs are indicated with their corresponding numbers) are displayed along the protein as dotted lines or shaded ranges. Numbers above each cross-linked subunit indicate TAF1 positions or range involved in the crosslinks, and the values in brackets indicate the number of distinct TAF1 cross-linked positions. Only crosslinks reported in at least two independent datasets are shown. The known structural domains of TAF1 are depicted. **b**, Heatmaps of TAF1 conservation (ConSurf), structural disorder (Metapredict), and pLDDT (AlphaFold structural prediction confidence) scores are represented. pLDDT > 90 are expected to be modelled to high accuracy, pLDDT between 70 and 90 are expected to be modelled well and pLDDT between 50 and 70 should be treated with caution. The scales of the different predictions scores are shown on the right. **c**, Full-length human TAF1 AlphaFold model. The initial structure was extended to better appreciate the different domains indicated along the protein. **d**-**e**-**f**, Structural models of the three main TAF1 anchor points (labelled A, B and C) in TFIID are shown in the insets. Model in (d) is the result of a AlphaFold prediction of the TAF1/TAF11/TAF13/TBP subcomplex. Distinct TAF1 domains are colored as in (a). Partner subunits are shown in shades of grey. TAF1 segments part of each anchor point are indicated. **g**, Schematic summary of the distinct interaction hotspots along the protein and the length of the intervening linker regions. TFIID lobe A and B are schematized by circles. TAND: TAF1 N-terminal domain; T6BM: TAF6 binding motif; RAPiD: RAP74 interaction domain; HMG: HMG-box domain; ZnK: zinc-knuckle domain; BD: bromodomain; HEAT: TAF6 HEAT repeat domain.

TAF1 modular organization is shown on the AlphaFold model of the full-length protein (**Fig. 5c**). A substantial fraction of the protein (∼48%) is predicted to be intrinsically disordered, including interdomain linker regions and the long acidic C-terminal tail (**Fig. 5b**). TAF1 contains two main well-structured regions: the TAF7 interaction domain (TAF7iD), which occupies the central portion of the protein, and two histone-reader bromodomains (BD) localized in tandem along the C-terminal tail (**Fig. 5a**). The TAF7iD – composed of the DUF3591 domain in concert with the RAP74 interaction domain (RAPiD) – tightly associates with TAF7 and binds downstream of core promoter DNA^18, 28^. Accordingly, these regions were modelled with high confidence by AlphaFold (**Fig. 5b**). Moreover, the scarcity of TAF1 intraprotein crosslinks outside the TAF7iD and the tandem-BDs is in accordance with the absence of other major structured domains along the protein (**Extended Data Fig. 5a**).

The three described crosslinking hotspots correspond to distinct anchor points (named here A-B-C) for specific TFIID submodules (**Fig. 5d-f**). The first TAF1 anchor point (A) would interact with TBP and the TAF11/TAF13 heterodimer. Indeed, the flexible TAF N-terminal domain (TAND) was shown to directly interact with and inhibit TBP DNA binding^29–31^. Additionally, removal of the human TAND abolished the co-translational recruitment of TBP to TAF1^24^. Concerning the TAF11/TAF13 heterodimer, the crosslinks with TAF1 are consistently found in all datasets. They map on TAF1 Lys249, lying within a conserved motif predicted with higher confidence and lower disorder scores with respect to the flanking regions (**Fig. 5a-c**). Modelling TAF1 with TBP and TAF11/TAF13 with AlphaFold resulted in a ternary complex with the expected positioning of TAF1 TAND into the concave surface of TBP. The putative TAF11/TAF13 interaction motif of TAF1 was folded laterally in a pocket formed by the HFDs subunits (**Fig. 5d**). Notably, TAF1 Lys249 position in the model is compatible with all the experimental crosslinks with TAF11/TAF13 (**Extended Data Fig. 5b**).

The second hotspot (B) is the anchor point of both lobes A and B with TAF1. It is constituted by three TAF1 stretches of conserved amino acids interspersed by loops of lower conservation, named TAF6-binding motifs (T6BMs, **Fig. 5a-c**). These motifs – recently resolved by cryo-EM^10^ – bridge the two copies of TAF6 HEAT domains, which in turn are connected to lobes A and B (**Fig. 5e**, see also **Fig. 1a**). The T6BMs occupy defined grooves and pockets across the pair of TAF6 HEAT domains at the center of TFIID (**Fig. 5e**). Modeling the entire TAF1 region containing the T6BMs allowed to map all crosslinking sites otherwise positioned in unresolved flexible loops (**Extended Data Fig. 5c**), in perfect agreement with the experimental structure (**Extended Data Fig. 5d-f**). Overall, the T6BMs constitute most of the interface anchoring together the two copies of TAF6 HEAT domains (**Extended Data Fig. 5g-h**). Note that the absence of crosslinked positions along the third T6BM (**Fig. 5a**) are due to the lack of lysine residues. Apart from TAF6, the crosslinks to other TAFs within this hotspot are likely driven by proximity rather than direct interactions.

The third and last hotspot (C) coincides with the TAF7iD (**Fig. 5f**). Besides the intricate fold adopted with TAF7, the DUF3591 loosely anchors the resulting TAF1/TAF7 globular domain to TAF2 (**Extended Data Fig. 5i**). Overall, structural and biochemical data support a scaffolding function of TAF1 within TFIID thanks to the modular organization of TAF1 domains (**Fig. 5g**). In this regard, TAF1 represents a flexible three-way anchor point that physically connects the three TFIID lobes through direct interactions with the two copies of TAF6, which in turn emanate into lobes A and B (**Fig. 5g**). One striking observation is that all mapped crosslinks reside in the N-terminal half of TAF1 (**Fig. 5a**), leaving the region downstream of RAPiD free from crosslinks (∼700 aa). This would potentially allow TFIID assembly on the N-terminal half of TAF1 before the protein is released from the ribosome.

### TAF1 depletion leads to an accumulation of TFIID building blocks in the cytoplasm

To investigate the role of TAF1 in the dynamics of cytoplasmic TFIID assembly, we perturbed the suggested TAF1-dependent assembly by siRNA-mediated TAF1 knockdown (KD). Subcellular fractionation experiments revealed an enrichment of the protein levels of several TFIID subunits in the cytoplasmic fraction upon TAF1 KD. Specifically, the cytoplasmic extract was substantially enriched for core-TFIID subunits (TAF4/5/6/12) when compared to the control-siRNA condition (**Fig. 6a**). Importantly, this cytoplasmic increase in protein levels of TAF4/5/6/12 was not visible in the nuclear fraction, suggesting a specific cytoplasmic accumulation of those subunits. Also, TAF13 and, to a lesser extent, TBP followed the same pattern. Instead, the levels of TAF8 and its partner TAF10 remained mostly unchanged. Notably, while the levels of TAF7 stayed constant in the cytoplasm, they were drastically reduced in the nuclear fraction, closely matching the depletion of TAF1. These observations show that TFIID subunits are differentially affected by TAF1 depletion. On the contrary, TAF4 and TAF7 KDs performed under the same conditions did not reproduce the effect elicited by TAF1 silencing, suggesting that the observed phenomenon is TAF1-specific (**Extended Data Fig. 6a**).

**Figure 6.**
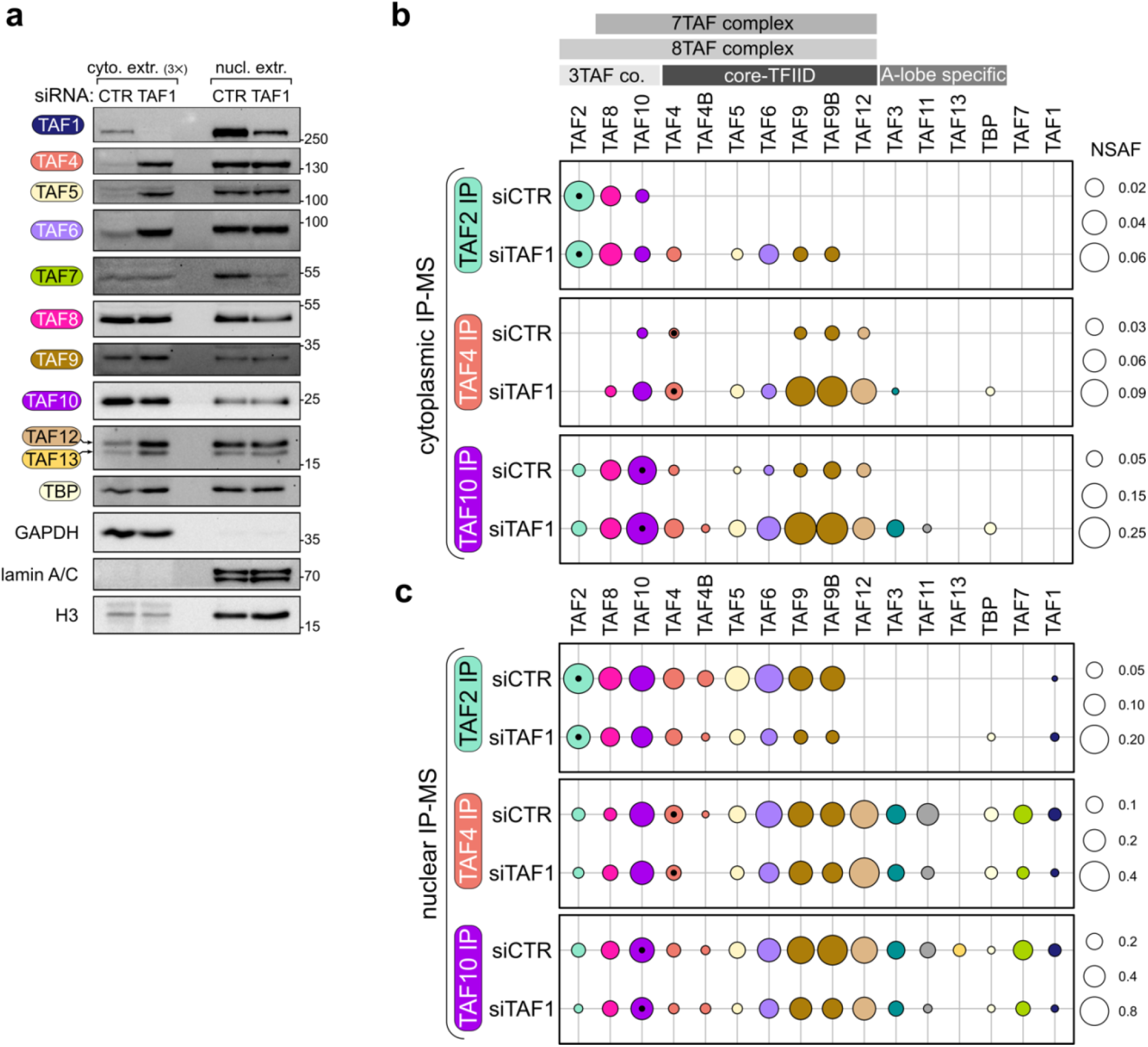
TAF1 depletion leads to an accumulation of TFIID building blocks in the cytoplasm. **a**, Subcellular fractionation of siRNA-transfected HeLa cells followed by western-blot analysis of endogenous TFIID subunits distribution. GAPDH, lamin A/C and histone H3 were used as loading controls. The amount of loaded cytoplasmic (cyto.) extract is three-times the amount of the nuclear (nucl.) extract counterpart. The positions of the molecular weight markers in kDa are indicated on the right. **b**, Immunoprecipitation (IP) of endogenous TFIID subunits coupled to label-free mass-spectrometry (MS) performed on cytoplasmic extracts of siRNA-transfected HeLa cells. Circle area represents the average NSAF value for each detected subunit in technical triplicates. The NSAF scales are indicated on the right. Distinct TFIID sub-complexes are depicted on the top. Subunits are color-coded and arranged according to the subcomplexes indicated on the top. Black dots in the circles identify the protein used as bait in each IP. **c**, Same as in (b) but the IPs were performed on nuclear extracts. CTR: non-targeting control siRNA. Co.: complex.

To address whether the cytoplasmic increase of a subset of TFIID subunits would correspond to an accumulation of specific TFIID building blocks in the cytoplasm, we analyzed endogenous cytoplasmic subcomplexes composition by performing IP-MS experiments upon TAF1 KD (**Fig. 6b** and **Extended Data Fig. 6b**). We selected IP-grade antibodies raised against a lobe C subunit (TAF2), a core TFIID subunit (TAF4) and a non-core TFIID subunit (TAF10). Note that we had to reduce the amount of input cells for the feasibility of siRNA transfection experiments coupled with IP-MS, at the cost of lower sensitivity. In good agreement with our findings, upon TAF1 KD, TFIID building blocks accumulated in the cytoplasm. In control condition, TAF2 was found associated with TAF8/TAF10 in the well-characterized 3TAF complex. However, following TAF1 KD, under these conditions core TFIID subunits joined the assembly to form the 8TAF complex. The enrichment of cytoplasmic core-TFIID was evidenced in all IPs. Notably, the lobe A-specific subunits, namely TAF3, TAF11 and TBP co-purified with TAF4 and TAF10, only following TAF1 depletion. We interpret these results as the cytoplasmic accumulation of different TFIID building blocks, including the 8TAF complex (B lobe + TAF2) and the A lobe complex, provoked by the impairment of the last, TAF1-dependent, assembly step before nuclear import.

Analyses of IP-MS on the nuclear fraction showed less dramatic rearrangements in subunits distribution, with an overall decrease in the abundance of all immunopurified TFIID subunits in TAF1 KD samples (**Fig. 6c** and **Extended Data Fig. 6c**). Overall, our observations together show that TAF1 is a major hub for the co-translational assembly of TFIID complex from preassembled building blocks and subsequent nuclear translocation.

## DISCUSSION

The self-assembly of large heterotypic multiprotein complexes in living cells poses major challenges to our understanding of cellular homeostasis. Here, we tackled the longstanding question of where and how the basal transcription factor TFIID assembles. To expand on our original findings, we comprehensively explored the landscape of co-translational assembly (Co-TA) events within TFIID, resulting in several novel pairs of subunits undergoing Co-TA (see **Table S1**). Most strikingly, we identified TAF1 as the central hub in the process.

### A hierarchical co-translational model for TFIID assembly

All our findings can be rationalized in a hierarchical model for TFIID assembly, which is stratified in three levels – or tiers – of assembly events (**Fig. 7**). The first tier includes early events along the pathway: these are the formation of protein pairs, mostly through the dimerization of HFD-containing subunits. We find it remarkable that all the HFD-pairs in TFIID assemble co-translationally, either directionally or symmetrically. This points at the histone-fold as a driver for Co-TA. The fact that several subunits used as bait in our cytoplasmic IP-MS data are not found as free proteins (**Fig. 4**) points either at a fast and efficient Co-TA with their partners, or to a degradation-driven removal of orphan subunits, although a combination of the two processes is also likely. Tier-1 also harbors interactions of non-HFD subunits, such as TAF2 and TAF5, which interact co-translationally with TAF8 and TAF6, respectively. Importantly, all these direct interacting pairs are structurally well characterized^10, 18^. The products of tier-1 assembly line are free early multisubunit intermediates, likely stabilized by the partners’ interactions. They are likely characterized by heterogenous half-lives as free molecular species, since some of them can be isolated in our steady state experiments (e.g. TAF11/TAF13), while others can only be detected as part of larger assemblies (e.g. TAF4/TAF12), yet some others are not detected at all (e.g. TAF3/TAF10) (**Fig. 4**). The products of tier-1 in turn access the second level of the assembly pathway by combining with each other in few – structurally constrained – steps. Importantly, assembly in tier-2 occurs post-translationally and leads to the buildup of larger assemblies recurrently found in our IP-MS experiments, such as the 8TAF complex and a partially assembled lobe A.

**Figure 7.**
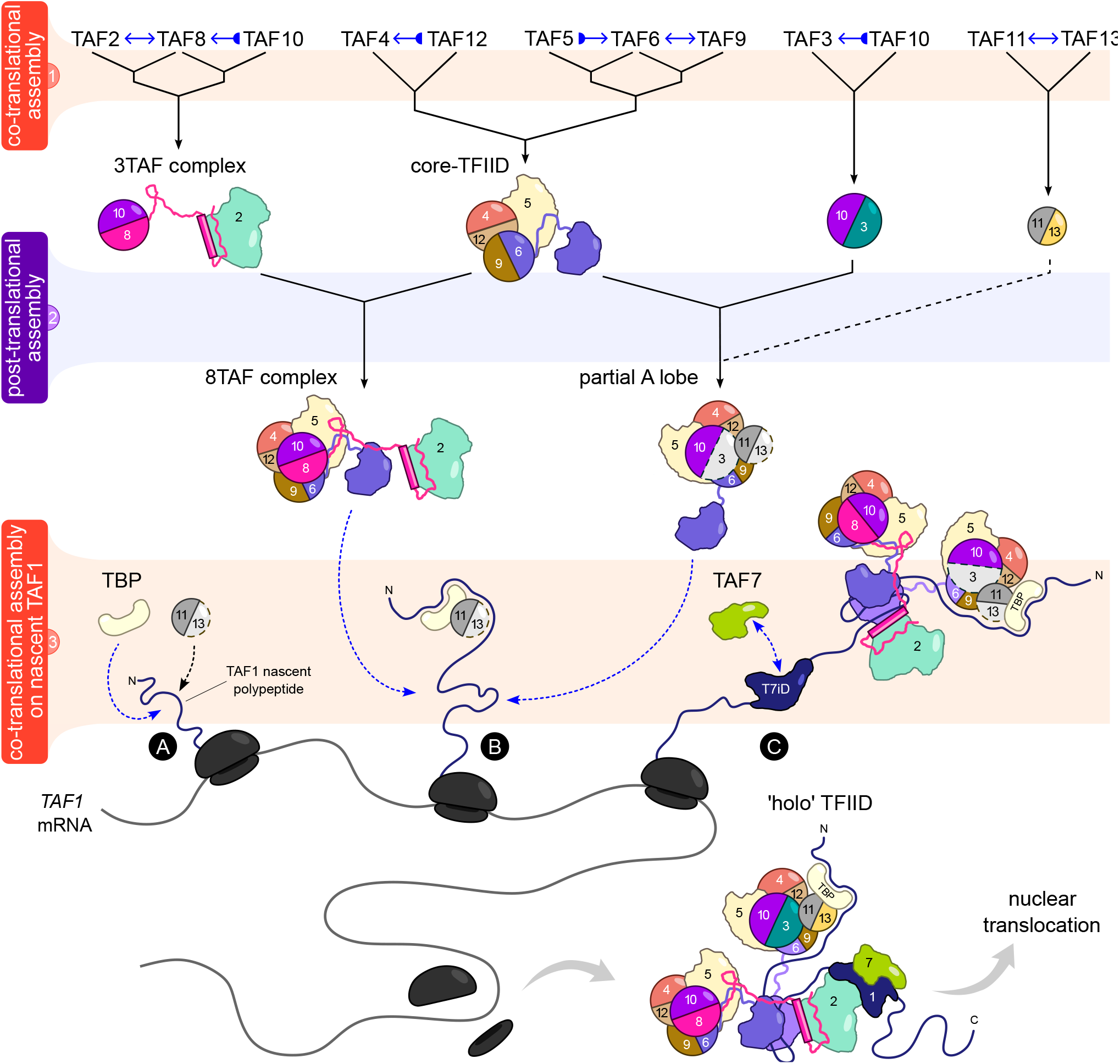
A co-translational hierarchical model for TFIID assembly. Scheme of the proposed cytoplasmic assembly model for TFIID that reconciles the experimental observations of the present work with previous structural and biochemical data. The assembly pathway can be subdivided in three tiers (colored and numbered horizontal stripes). Tier 3 represents the co-translational assembly of several TFIID building blocks on nascent TAF1 protein through three distinct interaction hotspots (labelled A, B and C), resulting in TFIID. Blue arrows with bases indicate directional Co-TA events, whereby double-headed blue arrows specify reciprocal Co-TA, as assessed by RIPs. For further details see Discussion.

In tier-3, the products of tier-2 finally converge and engage co-translationally with the nascent TAF1 polypeptide (**Fig. 7**). It is appealing to conceive a sequential N- to C-terminal order of assembly, whereby different TFIID building blocks are recruited by the distinct assembly domains of nascent TAF1 as the distinct interaction hotspots emerge from the ribosome channel. The first N-terminal anchor point (A) would interact with TBP, which engages with nascent TAF1 by binding the TAND domain^24^. Our systematic survey confirmed this Co-TA pair. TAF11/TAF13 dimer could also engage with nascent TAF1 at anchor point (A), forming a ternary complex along with TBP (**Fig. 5d**). Biochemically, a recombinant complex formed by TAF1/TBP/TAF11/TAF13 and TAF7 can be readily purified^32^ and the direct interaction between TAF1 and TAF11/TAF13 is supported by crosslinking experiments and structural modelling (**Fig. 5c-d**, **Extended Data Fig. 5b**). On the other hand, TAF11/TAF13 did not score positive for *TAF1* mRNA in our systematic RIP approach, opening the possibility of post-translational engagement, or weaker interactions unable to withstand the stringent RIP conditions. Alternatively, a preassembled TBP/TAF11/TAF13 sub-complex may interact with the N-terminal end of nascent TAF1.

The second interaction anchor point (B) – the T6BMs – would interact with two copies of TAF6 HEAT domains, effectively bringing together lobe A and lobe B (**Fig. 7**, **Fig. 5g**). Interestingly, a single protein – TAF1 – evolved distinct binding motifs able to recognize corresponding identical surfaces from each the two copies of TAF6 within TFIID (**Extended Data Fig. 5h**). The third anchor point (C) recruits TAF7, which is known to interact with the TAF1 central domain (DUF3591/RAPiD) (**Fig. 5g**). It is noteworthy to point out that in our RIP experiments TAF7 and TAF1 scored reciprocally positive, opening the possibility of a simultaneous co-translational interaction between the two. Such an ordered addition would entail a remarkable degree of coordination, potentially reinforced by binding cooperativity phenomena among the modules as they join the growing assembly. Yet, in our imaging data we detected TAF7-positive *TAF1* RNA spots lacking TBP signal and *vice versa*, hinting at a potential independent binding mode also (**Extended Data Fig. 3**).

Upon completion of TAF1 protein synthesis, a fully assembled TFIID is released and readily translocated in the nucleus. We point out that a subset of subunits scored negative for TAF1 mRNA in our RIP assays: these include TAF3, TAF9, TAF11 and TAF13. Therefore, it is possible that they join the complex post-translationally or through their interaction partners. The benefits of a hierarchical co-translational assembly have been recently theorized in the framework of yeast nuclear pore assembly^9^. The intriguing model proposed by the authors substantially applies also to our findings, where Co-TA is pervasively exploited for the hierarchical assembly of TFIID in the cytoplasm of mammalian cells.

All our present data is in agreement with the published TAFs interactions and the cryo-EM TFIID structures^10, 23^. As a consequence, our assembly pathway is also in agreement with previously published descriptions of partial TFIID assemblies ^27, 33^. Based on recent bioinformatic analyses, during evolution proteins that assemble co-translationally have sustained large N-terminal interfaces in order to promote co-translational subunit recruitment^34^. In agreement, out of the eight larger subunits of TFIID that participate in Co-TA as nascent polypeptides (TAF1, 2, 3, 4, 6, 7, 8, 9), seven have their interaction domains located in the N-terminal of the given subunit (TAF1, 2, 3, 6, 7, 8, 9).

### TAF1: a ‘driver’ and limiting factor along the assembly line

*TAF1* mRNA was enriched in the majority of our RIPs (**Fig. 1**, **Table S1**) and it was found in physical proximity with several TFIID subunits in the cytoplasm (**Fig. 3**). As suggested by our cytoplasmic IP-MS data (**Fig. 4**), in this compartment the levels of TAF1 protein seem to be limiting with respect to TFIID building blocks. Notably, TAF7 IP enriched the whole spectrum of TFIID subunits, TAF1 included, albeit at very low levels (**Fig. 4e** and **Extended Data Fig 4b**). TAF7 also showed the highest levels of co-localization with *TAF1* mRNA (**Fig. 3d**), pointing at a remarkable Co-TA efficiency between TAF1 and TAF7. The higher assembly efficiency is consistent with the detection of a fully assembled complex in TAF7 cytoplasmic IP. It is worth to emphasize that the interaction interface between TAF1 and TAF7 is remarkably intricate, with deeply intertwined β-strands from each protein contributing to a common β-barrel^28^. In fact, it would be conceivable that such an interface would only form concomitantly with folding during protein synthesis, imposing a structural constraint solved by Co-TA. Curiously, TAF7 levels decreased proportionally with TAF1 depletion in the nucleus, hinting at a partner stabilization effect (**Fig. 6a**), similar to the one observed between TAF10 and TAF8^24^, or impaired nuclear import. Importantly however, TAF1 depletion led to the accumulation of several TFIID building blocks in the cytoplasm, revealing a key role of this subunit in driving complex assembly and consequent relocation in the nucleus (**Fig. 6**). We propose that nascent TAF1 nucleates the late steps of TFIID assembly in the cytoplasm by tethering together different sub-modules of the complex, and – once released from the ribosomes – the whole assembly efficiently shuttles in the nucleus (**Fig. 7**). This process may act as a quality checkpoint before nuclear import. In agreement, both in yeast and in metazoans *TAF1* is an essential gene^35–37^.

A central role of a single nascent subunit for the co-translational assembly of protein complexes has been demonstrated for the COMPASS histone methyltransferase in budding yeast, where a specific sub-complex is directly assembled on nascent Set1 protein, stabilizing the latter from degradation^38^. In this context Set1 behaved as a co-translational ‘driver’ subunit, simultaneously promoting complex assembly and limiting its abundance. Although not as thoroughly dissected, other examples of central subunits potentially working as co-translational drivers in the context of complex assemblies were evidenced in fission yeast^4^ and supported by structural analyses^39^. We argue that an equivalent process in mammalian cells is led by TAF1 as the driver subunit for TFIID co-translational assembly, which culminates with the tethering of distinct building blocks on TAF1 nascent polypeptide. In this regard, *TAF1* mRNA offers the longest coding sequence (CDS) among TFIID components, implying a prolonged timeframe for co-translational binding events to occur. By taking into account an estimate of average translation speed of ∼5.6 codons per second in mammalian cells^40^, translating *TAF1* CDS would take ∼ 5.6 min. The last assembly domain along TAF1 completely emerges from the ribosome around position 1240, granting an additional window of time of about 1.9 min to ultimate Co-TA before ribosome release. In this regard, the analysis of ribosome profiling (Ribo-seq) merged datasets showed a wide region of sparse ribosome-protected fragments, encompassing all three T6BMs and extending inside the DUF3591 domain-encoding region (**Extended Data Fig. 7**). The low signal in the T6BMs region hints at fast elongation rates, which would rapidly expose all three T6BMs for the co-translational recruitment of the respective TFIID building-blocks. Downstream to this region, translation significantly slows down, as suggested by the higher ribosome occupancy. This seems to occur once the synthesis of the heavily structured DUF3591 central domain has started. In this view, contrary to the simple short linear motifs represented by the T6BMs, the co-translational folding of DUF3591 domain with TAF7 might benefit from a slower translation pace. This would also buy time to establish productive co-translational interactions with the anchor points located upstream.

### Open questions

Our findings reveal an unprecedented mechanism for TFIID biogenesis, answering longstanding questions and opening new ones. One of remarkable importance is how efficiently does Co-TA occur. A prerequisite for co-translational interactions is an actively translated mRNA. Our imaging approach detected nascent TAF1 protein on roughly half of the correspondent cytosolic messengers (**Fig. 2**). By using this observation as a proxy for the proportion of actively translated *TAF1* mRNAs, the observed frequency of Co-TA events for the other probed subunits (**Fig. 3**) would rather be underestimated. In the future, the observation of co-translational binding events in living cells might offer a quantitative dimension to this field.

A second point is whether Co-TA in TFIID is an efficient option for complex assembly or rather an obligate path. The same question can be extended to other molecular complexes as well. Co-TA might be the sole opportunity for assembly domains characterized by structural constraints, such as the TAF1/TAF7 interface. Instead, interactions mediated by classical binding pockets, extended surfaces or short linear motifs can rely also on post-translational assembly. However, co-translational interactions have the advantage of abolishing partially unfolded/unstable intermediates by kinetically anticipating their complexed state. Although it seems reasonable to hypothesize that natural selection promoted molecular features favoring co-translational interactions, it has proven hard to disentangle co-translational from post-translational assembly experimentally, since both mechanisms ultimately depend on protein synthesis. As yeast Tra1, a large pseudokinase and a subunit of SAGA and NuA4 complexes, uses chaperone-mediated assembly processes^41^, other assembly mechanisms likely play a role in multi-subunit complex assembly pathways.

Third, our data open new questions on the nuclear import mechanism adopted by TFIID or its building blocks. The observation that a defined set of subcomplexes accumulates in the cytoplasm upon TAF1 depletion opens the possibility of distinct entry routes to the nucleus. A fully assembled – TAF1-containing – complex could represent the most efficiently translocated molecular species, with several subcomplexes relying on TAF1 for nuclear import. Conversely, other building blocks might access the nucleus autonomously, eventually less efficiently though, as demonstrated in the past for the TAF2/TAF8/TAF10 module^21, 42^. Interestingly, the interrogation of a systematic interactome survey of the major nuclear transport receptors on human cells by BioID^43^ showed that all the detected TFIID subunits shared the same import systems, mainly the α-importins IMA1 and IMA5. This is consistent with the idea that TFIID is transported across the nuclear envelope as a pre-assembled entity. Intriguingly, in this study TAF1 was one of the main biotinylated TFIID subunit, suggesting that TAF1 can directly interact with the nuclear transport receptors and potentially drive nuclear import.

The picture that emerges from the present study provides a novel understanding of the complex series of steps underlying the assembly mechanism of the basal transcription factor TFIID. We envision that the principles of hierarchical co-translational assembly could apply to the biogenesis of most large heteromeric multiprotein complexes in living cells.

## METHODS

### Cell culture

Human HeLa cells (CCL-2; ATCC) were obtained from the IGBMC cell culture facility and cultured in DMEM (4.5 g/L glucose) supplemented with 10% fetal calf serum (Dutscher, S1810) and 100 U/mL penicillin 100 μg/mL streptomycin (Invitrogen, 15140-130). E14 mouse embryonic stem cells (mESCs, ES Parental cell line E14Tg2a.4, Mutant Mouse Resource and Research Center) were obtained from the IGBMC cell culture facility and cultured on gelatinized plates in feeder-free conditions in KnockOut DMEM (Gibco) supplemented with 20 mM L-glutamine, pen/strep, 100 µM non-essential amino acids, 100 µM β-mercaptoethanol, N-2 supplement, B-27 supplement, 1000 U/mL LIF (Millipore), 15% ESQ FBS (Gibco) and 2i (3 µM CHIR99021, 1 µM PD0325901, Axon MedChem). Cells were grown at 37°C in a humidified, 5% CO2 incubator.

### GFP-fusion cell lines generation

GFP-TAFs fusion cell lines used in this study were described in^26^. Briefly, the coding sequences for the human TFIID subunits (TAF1, TAF2, TAF3, TAF4, TAF5, TAF6, TAF7, TAF8, TAF9, TAF10, TAF11, TAF12, TAF13 and TBP) were obtained by PCR using the appropriate cDNA clone and gene-specific primers flanked by attB sites followed by BP-mediated GATEWAY recombination into pDONR221 according to manufacturer’s instructions (Invitrogen). The cloned sequence was verified by sequencing and it was transferred to the pcDNA5-FRT-TO-N-GFP Gateway destination vector by LR recombination according to the manufacturer’s protocol (Invitrogen). HeLa Flp-In/T-REx cells, which contain a single FRT site and express the Tet repressor^25^, were grown in DMEM, 4.5 g/L glucose (Gibco), supplemented with 10% v/v fetal calf serum (Gibco). All the GFP-fusion destination vectors were co-transfected with a pOG44 plasmid that encodes the Flp recombinase into HeLa Flp-In/T-REx cells using polyethyleneimine (PEI) to generate stable doxycycline-inducible expression cell lines. Recombined cells were selected with 5 μg/mL blasticidin S (InvivoGen) and 250 μg/mL hygromycin B (Roche Diagnostics) 48 h after PEI transfection. Cells were maintained in DMEM supplemented with 10% Tet-free fetal calf serum (Pan Biotech, P30-3602), blasticidin S, hygromycin B and pen/strep.

### RNA-immunoprecipitation (RIP) against endogenous TFIID subunits

Polysome extract preparation and RIPs were performed essentially as described in^24^. HeLa cells grown on 15 cm plates (∼90% confluent) were treated either with 100 μg/mL cycloheximide (CHX, Merck, C1988) for 15 min or with 50 μg/mL puromycin (Puro, Invivogen, ant-pr-1) for 30 min in the incubator at 37°C. Plates were placed on ice and cells were washed twice with ice-cold PBS and scraped in 2 mL lysis buffer [20 mM HEPES-KOH pH 7.5, 150 mM KCl, 10 mM MgCl2, 0.1% NP-40, 1× PIC (cOmplete EDTA-free protease inhibitor cocktail, Roche, 11873580001), 0.5 mM DTT (ThermoScientific, R0862), 40 U/mL RNasin Ribonuclease Inhibitor (Promega, N2511)] supplemented either with CHX or Puro. Cell suspension was homogenized with 10 Dounce strokes using a B-type pestle on ice. Lysates were incubated 15 min on ice and cleared by centrifugation at 17000 × g. The supernatant represents the polysome extract.

1.2 mg protein G Dynabeads (Invitrogen, 10004D) were used for each immunoprecipitation (IP). The antibodies employed are listed in **Table S3**. Dynabeads were washed twice in buffer IP100 (25 mM Tris-HCl pH 8.0, 100 mM KCl, 5 mM MgCl2, 10% glycerol, 0.1% NP-40). Each antibody (5-10 μg/IP) was coupled to Dynabeads in 100 μL buffer IP100 for 1 hour at room temperature (RT) in agitation. Mock IPs were performed using mouse or rabbit IgG. Antibody-coupled Dynabeads were washed twice in buffer IP500 (25 mM Tris-HCl pH 8.0, 500 mM KCl, 5 mM MgCl2, 10% glycerol, 0.1% NP-40) and three times in buffer IP100. 1 mL of polysome extract (equivalent ∼10^7^ cells) was used as input for each IP. A 10% equivalent volume of the input was kept at 4°C for input-normalization. IP reactions were incubated in rotation at 4°C overnight. The next day Dynabeads were washed four times with 0.5 mL high salt was buffer (25 mM HEPES-KOH pH 7.5, 350 mM KCl, 10 mM MgCl2, 0.02% NP-40, 1× PIC, 0.5 mM DTT, 40 U/mL RNasin Ribonuclease Inhibitor) supplemented either with CHX or Puro. RNA from the resulting immunopurified material was extracted using NucleoSpin RNA XS kit (Macherey-Nagel, 740902) in 100 μL RA1 lysis buffer and purified according to the manufacturer’s protocol. The input sample was extracted and processed in parallel with the IPs.

### GFP-fusion RIP

GFP-RIPs using inducible cell lines were performed as described for endogenous RIPs, with the following modifications. The day of the experiment, the expression of the GFP-tagged TFIID subunit was induced by addition of 1 μg/mL doxycycline (Dox) for 2 hours. For GFP-TAF3 and GFP-TAF10 cell lines Dox treatment was omitted due to their leaky basal expression. Cells were treated with CHX, lysed and polysome extracts prepared as described in the previous section. GFP-IPs were carried out using 40 μL GFP-Trap Agarose beads (ChromoTek, gta-20). Mock IPs were carried out using an equivalent volume of protein G Sepharose beads. Beads were incubated with polysome extracts for 4 hours at 4°C, washed and RNA purified as described in the previous section.

### RT-qPCR

Reverse transcription reaction was performed using SuperScript IV First-Strand Synthesis System (Invitrogen, 18091050) and random hexamers according to manufacturer instructions. The resulting cDNA was diluted 1:10. Quantitative PCR was performed with two or three technical replicas using 2 μL cDNA, primers listed in **Table S3**, and LightCycler 480 SYBR Green I Master (Roche, 04887352001), in a LightCycler 480 instrument (Roche). Input % normalization for RIP samples was performed by applying the formula 100 × 2 ^ [(Ctinput – 6.644) – CtRIP]. Fold-enrichment normalization was performed by dividing RIP Input % by mock Input %.

### Immunofluorescence (IF) coupled to single-molecule inexpensive RNA FISH (smiFISH)

RNA detection was performed through smiFISH^44^. Primary probe sets (24 single oligonucleotides) against target coding sequence were designed using Oligostan in R as described in the software documentation^44^. Probes sequences are reported in **Table S3**. Primary probes were synthesized by Integrated DNA Technologies (IDT) in plate format, dissolved in TE buffer at 100 μM. 5’ and 3’ Cy3-labelled secondary probe (FLAP) was synthesized by IDT and purified by HPLC. Primary probes were mixed in an equimolar solution in TE at 0.83 μM per probe. To prepare a 50× smiFISH composite probes mix, 4 μL primary probes mix were mixed with 2 μL 100 μM secondary probe solution in 20 μL final reaction volume in 100 mM NaCl, 50 mM Tris-HCl pH 8.0, 10 mM MgCl2. The annealing reaction was performed in a thermocycler with the following conditions: 3 min at 85°C, 3 min at 65°C, 5 min at 25°C. 50× smiFISH probes mix was stored at –20°C. The day before the experiment, HeLa cells were seeded on coverslips (No. 1.5H, Marienfeld, 630-2000) in a 12-well plate (0.2 × 106 cells/well). The day after, cells were treated either with 100 μg/mL CHX for 15 min or with 50 μg/mL Puro for 30 min in the incubator at 37°C. Then, cells were directly processed for immunofluorescence. All buffer solutions were filtered (0.22 μm filter). Cells were washed twice with PBS (containing CHX for cells treated with it) and fixed with 4% paraformaldehyde (Electron Microscopy Sciences, 15710) in PBS for 10 min at RT. Cells were washed twice with PBS and incubated for 10 min at RT in blocking/permeabilization solution (BPS) [1× PBS, 1% BSA (MP, 160069), 0.1% Triton-X100 (Merck, T8787), 2 mM vanadyl ribonucleoside complexes (VRC, Merck, R3380)]. Cells were incubated for 2 hours at RT with the following primary antibodies diluted in BPS: TAF1 (1:1000, rabbit pAb, Abcam, ab188427), TAF4 (3 μg/mL, mouse mAb, 32TA 2B9), TAF7 (1:250, rabbit pAb, #3475), TAF10 (3 μg/mL, mouse mAb, 6TA 2B11), TBP (2 μg/mL, mouse mAb, 3TF1 3G3) or SUPT7L (rabbit pAb, Bethyl, A302-803A). A secondary-only control sample was incubated with BPS devoid of primary antibody. After three 5 min PBS washes, cells were incubated for 1 hour at RT (light-protected) with AF488-conjiugated secondary antibodies diluted 1:3000 in BPS (goat anti-mouse IgG, A11001 or goat anti-rabbit IgG, A11008, Life Technologies). For dual-color IF, we also used AF(Plus)647-conjiugated secondary antibody (goat anti-mouse IgG, A32728). After three 5 min PBS washes, a second fixation step was performed with 4% paraformaldehyde in PBS for 10 min at RT. Cells were washed twice with PBS and equilibrated in hybridization buffer [2× SSC buffer, 10% formamide (Merck, F9037)] for at least 10 min at RT. An equivalent volume of the following mixes was prepared: Mix1 [2× smiFISH probes mix, 2× SSC buffer, 30% formamide, 0.68 mg/mL *E. coli* tRNA (Roche, 10109541001)] and Mix2 [0.4 mg/mL BSA (NEB, B9000S), 4 mM VRC, 21.6% dextran sulfate (Merck, D8906)]. Mix1 and Mix2 were combined 1:1 and thoroughly mixed by vortexing. 45 μL of the resulting solution were applied on the surface of a 10 cm plastic dish that served as hybridization chamber. Each coverslip was applied upside-down on the smiFISH mix drop. A hydration chamber (a 3.5 cm plate filled with hybridization buffer) was included. The hybridization chamber was sealed with parafilm and incubated overnight at 37°C, light protected. The day after, each coverslip was washed twice at 37°C for 30 min in 2 mL hybridization buffer. 0.5 μg/mL DAPI (Merck, MBD0015) was included in the second wash for nuclear counterstain. After two PBS washes, coverslips were mounted with 5 μL Vectashield (Vector Laboratories, H-1000) and sealed with nail polish.

### Confocal microscopy and image processing

Cells processed for immunofluorescence/smFISH were imaged using spinning disk confocal microscopy on an inverted Leica DMi8 equipped with a CSU-W1 confocal scanner unit (Yokogawa), with a 1.4 NA 63× oil-objective (HCX PL APO lambda blue) and an ORCA-Flash4.0 camera (Hamamatsu). DAPI, AF488 (IF) and Cy3 (smFISH) were excited using a 405 nm (20% laser power), 488 nm (70%) and 561 nm (70%) laser lines, respectively. For dual color IF experiments, AF(Plus)647 was excited using the 642 nm laser line. 3D image acquisition was managed using MetaMorph software (Molecular Devices). 2048×2048 pixels images (16-bit) were acquired with a xy pixel size of 0.103 μm and a z step size of 0.3 μm (∼30-40 optical slices). Multichannel acquisition was performed at each z-plane. Multicolor fluorescent beads (TetraSpeck Fluorescent Microspheres, Invitrogen, T14792) were imaged alongside the samples. Chromatic shift registration was performed with Chromagnon^45^ using the fluorescent beads hyperstack as reference. Image channels were split and maximum intensity projections (MIPs) were generated in Fiji^46^ using a macro. smFISH RNA spots were detected and counted using RS-FISH Fiji plugin^47^ on MIPs. Briefly, anisotropy coefficient calculation was performed on a smFISH z-stack image and spot detection on MIPs was performed in ‘advanced mode’ (no RANSAC, compute min/max intensity from image, use anisotropy coefficient for DoG, add detections to ROI-Manager, mean background subtraction, Sigma = 1.25, DoG and intensity thresholds were manually adjusted). All detected RNA spots were saved as ROI selections and used to create an RNA spots label map image (each spot is identified as a pixel with a distinct value) using a custom Fiji macro. A CellProfiler^48^ pipeline was used to segment cells and allocate/count cytoplasmic RNA spots. Briefly, DAPI images were used to identify nuclei as primary objects using minimum cross-entropy thresholding method, smFISH background fluorescence was used to identify cell boundaries as secondary objects and cytoplasmic regions were derived by subtracting nuclei from cells. The ‘RelateObjects’ function was used to assign each RNA spot to the mother object cytoplasm. Total number of cytoplasmic RNA spots per image was computed. To count the number of cytoplasmic RNA spots per cell, cells touching image border were excluded. The detection of cytoplasmic RNA spots (smFISH) co-localizing with protein spots (IF) was performed manually on chromatic shift-corrected multichannel z-stack images. To avoid operator bias in image annotation, image files were randomized using a custom Fiji macro script before the analysis. The position of cytoplasmic RNA spots was used as reference to check for the presence of resolution-limited particles in the IF channel, distinct from the background and overlapping in xyz with the RNA spots. The position of each positive co-localization event was recorded in ROI manager. To account for RNA abundance, the number of RNA spots co-localized with protein spots was normalized to the total number of cytoplasmic RNA spots per image and expressed as a fraction. If not specified otherwise, images shown in the main figures correspond to representative subsets of single optical planes from chromatic shift-corrected confocal images. Brightness and contrast adjustments were applied on the entire image in Fiji to facilitate the visualization, without background clipping.

### siRNA transfection

Control (siCTR) and TAF1 siRNAs were purchased from Horizon (ON-TARGETplus Non-targeting Control Pool D-001810-10-05, ON-TARGETplus Human TAF1 siRNA SMARTpool L- 005041-00-0010) and resuspended in nuclease-free H2O. For large-scale transfections, 2.5 ×10^6^ HeLa cells were seeded in 10 cm plates. The day after, cells were transfected using Lipofectamine 2000 (Invitrogen, 11668019) using a low-volume transfection protocol. In brief, after medium removal, cells were treated with 17.5 μL Lipofectamine 2000 diluted in 2.8 mL Opti-MEM (Gibco, 31985062) for 15 min at 37°C. Then, 56 pmol of siRNA diluted in 0.7 mL Opti-MEM were added dropwise to the cells and gently mixed, achieving 16 nM final siRNA concentration. After ∼5 hours incubation at 37°C, the transfection mix was replaced with prewarmed complete DMEM. Cells were harvested 48 hours post-transfection.

### Western blot

Samples were loaded on SDS-PAGE gels added with 0.5% 2,2,2-trichloroethanol (TCE, Sigma-Aldrich) for stain-free protein detection^49^. The gel was activated for one minute with UV and the proteins were transferred to a nitrocellulose membrane following standard procedures. Specific proteins were probed with the primary antibodies listed in **Table S3** and HRP-conjugated secondary antibodies. To reprobe the membrane with an antibody raised in a different species, the previous secondary antibody was inactivated with 10% acetic acid according to^50^.

### Subcellular fractionation

Adherent cells were washed with cold PBS twice and harvested by scraping on ice. Cell suspension was centrifuged at 400 ×g for 5 min at 4°C and the pellet was resuspended in 4 packed cell volumes (PCV) of hypotonic buffer (50 mM Tris-HCl pH 8.0, 1 mM EDTA, 1 mM DTT, 1× PIC). After 30 min incubation on ice, cells were lysed with 10 hits of Dounce homogenizer and centrifuged at 2300 ×g for 10 min at 4°C. The supernatant was saved as cytoplasmic extract. Nuclei were washed once in hypotonic buffer and resuspended in 3.5 PCV hypertonic buffer (50 mM Tris-HCl pH 8.0, 0.5 mM EDTA, 500 mM NaCl, 25% glycerol, 1 mM DTT, 1× PIC). Nuclei were lysed with 20 hits of Dounce homogenizer, incubated in agitation for 30 min at 4°C and centrifuged at 19000 ×g for 30 min at 4°C. The supernatant was saved as nuclear extract. Cytoplasmic and nuclear extracts were dialyzed against 25 mM Tris-HCl pH 8.0, 5 mM MgCl2, 100 mM KCl, 10% glycerol, 0.5 mM DTT, 1× PIC at 4°C using DiaEasy dialyzers (BioVision K1013-10) and protein concentration was measured using Bradford assay (BioRad, 5000006).

### Immunoprecipitation coupled to LC-MS/MS analysis

Specific and mock (anti-GST) antibodies were coupled either with 200 μL protein G-Sepharose (large scale IPs, **Fig. 4** and **Extended Data Fig. 4**) or with 2.7 mg protein G Dynabeads (medium scale IPs, **Fig. 6**) in IP100 buffer (25 mM Tris-HCl pH 8.0, 5 mM MgCl2, 100 mM KCl, 10% glycerol, 0.1% NP-40, 0.5 mM DTT, 1× PIC) in agitation for 1 hour at RT. Antibody-coupled beads were washed twice in IP500 buffer (25 mM Tris-HCl pH 8.0, 500 mM KCl, 5 mM MgCl2, 10% glycerol, 0.1% NP-40, 0.5 mM DTT, 1× PIC) and three times in buffer IP100. Antibody-coupled beads were incubated with cytoplasmic (3-30 mg, medium-large scale IPs) or nuclear (1-10 mg) extracts overnight at 4°C. The day after, beads were washed twice with IP500 for 5 min at 4°C and three times with IP100. Immunopurified proteins were eluted in 0.1 M glycine pH 2.7 and immediately buffered with 1 M Tris-HCl pH 8.0. Eluates were precipitated with TCA (Merck, T0699) overnight at 4°C and centrifuged at 14000 ×g for 30 min at 4°C. Protein pellets were washed twice with cold acetone and centrifuged at 14000 ×g for 10 min at 4 °C. Pellets were denatured with 8 M urea (Merck, U0631) in 0.1 M Tris-HCl, reduced with 5 mM TCEP for 30 min and alkylated with 10 mM iodoacetamide (Merck, I1149) for 30 min, light-protected. Both reduction and alkylation were performed at RT and in agitation. Double digestion was performed with endoproteinase Lys-C (Wako, 125-05061) at a 1:100 ratio (enzyme:protein) in 8 M urea for 4 hours, followed by an overnight modified trypsin digestion (Promega, V5113) at a 1:100 ratio in 2 M urea for 12 hours.

Samples were analyzed using an Ultimate 3000 nano-RSLC coupled in line, via a nano-electrospray ionization source, with the LTQ-Orbitrap ELITE mass spectrometer (Thermo Fisher Scientific) or with the Orbitrap Exploris 480 mass-spectrometer (Thermo Fisher Scientific) equipped with a FAIMS (high Field Asymmetric Ion Mobility Spectrometry) module. Peptide mixtures were injected in 0.1% TFA on a C18 Acclaim PepMap100 trap-column (75 µm ID x 2 cm, 3 µm, 100 Å, Thermo Fisher Scientific) for 3 min at 5 µL/min with 2% ACN, 0.1% FA in H2O and then separated on a C18 Acclaim PepMap100 nano-column (75 µm ID x 50 cm, 2.6 µm, 150 Å, Thermo Fisher Scientific) at 300 nL/min, at 40°C with a 90 min linear gradient from 5% to 30% buffer B (A: 0.1% FA in H2O / B: 80% ACN, 0.1% FA in H2O), regeneration at 5% B. Spray voltage were set to 2.1 kV and heated capillary temperature at 280°C. For Orbitrap Elite, the mass spectrometer was operated in positive ionization mode, in data-dependent mode with survey scans from m/z 350-1500 acquired in the Orbitrap at a resolution of 120000 at m/z 400. The 20 most intense peaks from survey scans were selected for further fragmentation in the Linear Ion Trap with an isolation window of 2.0 Da and were fragmented by CID with normalized collision energy of 35% (TOP20CID method). Unassigned and single charged states were excluded from fragmentation. The Ion Target Value for the survey scans (in the Orbitrap) and the MS2 mode (in the Linear Ion Trap) were set to 1E6 and 5E3 respectively and the maximum injection time was set to 100 ms for both scan modes. Dynamic exclusion was set to 20 s after one repeat count with mass width at ± 10 ppm. For Orbitrap Exploris 480 MS associated with the FAIMS module, a combination of two Compensation Voltage (CV), −40 V and −55 V, was chosen with a cycle time of 1 s for each. For the full MS1 in DDA mode, the resolution was set to 60000 at m/z 200 and with a mass range set to 350-1400. The full MS AGC target was 300% with an IT set to Auto mode. For the fragment spectra in MS2, AGC target value was 100% (Standard) with a resolution of 30000 and the maximum Injection Time set to Auto mode. Intensity threshold was set at 1E4. Isolation width was set at 2 m/z and normalized collision energy was set at 30%. All spectra were acquired in centroid mode using positive polarity. Default settings were used for FAIMS with voltages applied as described previously, and with a total carrier gas flow set to 4.2 L/min.

### Mass spectrometry data analysis

Proteins were identified by database searching using SequestHT (Thermo Fisher Scientific) with Proteome Discoverer 2.4 software (PD2.4, Thermo Fisher Scientific) on human FASTA database downloaded from UniProt (reviewed, release 2021_06_03, 20380 entries, https://www.uniprot.org/). Precursor and fragment mass tolerances were set at 7 ppm and 0.6 Da respectively, and up to 2 missed cleavages were allowed. For the data acquired on the Orbitrap Exploris 480, the software Proteome Discoverer 2.5 version was used with a human fasta database from UniProt (reviewed, release 2022_02_21, 20291 entries). Precursor and fragment mass tolerances were set at 10 ppm and 0.02 Da respectively, and up to 2 missed cleavages were allowed. For all the data, Oxidation (M, +15.995 Da) was set as variable modification, and Carbamidomethylation (C, + 57.021 Da) as fixed modification. Peptides and proteins were filtered with a false discovery rate (FDR) at 1%. Label-free quantification was based on the extracted ion chromatography intensity of the peptides. All samples were measured in technical triplicates. The measured extracted ion chromatogram (XIC) intensities were normalized based on median intensities of the entire dataset to correct minor loading differences. For statistical tests and enrichment calculations, not detectable intensity values were treated with an imputation method, where the missing values were replaced by random values similar to the 10% of the lowest intensity values present in the entire dataset. Unpaired two tailed T-test, assuming equal variance, were performed on obtained log2 XIC intensities. Normalized spectral abundance factors (NSAF) were calculated for each protein as described earlier^51^. To obtain spectral abundance factors (SAF), spectral counts identifying a protein were divided by the protein length. To calculate NSAF values, the SAF values of each protein were divided by the sum of SAF values of all detected proteins in each run. All raw LC-MS/MS data have been deposited to the ProteomeXchange via the PRIDE repository with identifier PXD036358.

### Crosslinking-MS metanalysis, protein sequence analysis and modelling

For the metanalysis on the available crosslinking-MS experiments performed on human TFIID, we retrieved and combined the curated datasets from Patel et al., 2018^18^(one dataset, apo-TFIID), Scheer et al., 2021^19^(one dataset, apo-TFIID) and Chen et al., 2021^10^(five datasets of TFIID incorporated in preinitiation complex variants: cPICscp, cPICpuma, mPICscp, hPICscp, p53hPIChdm2), for a total of seven datasets. We only included intra and interprotein crosslinks involving TAF1 and found in at least two different datasets. If a crosslink was only present among the Chen et al. datasets, it was considered only if it scored as significant in more than one dataset (probability score <0.05). The resulting subset of common TAF1 crosslinks is reported in **Table S2**. TAF1 conservation and structural disorder prediction were computed using ConSurf^52^ and Metapredict^53^, respectively. TAF1 full-length model corresponding to the UniProt entry P21675 was downloaded from AlphaFold Protein Structure Database (https://alphafold.ebi.ac.uk/). For visual clarity in **Fig. 5c**, the model backbone was manually extended at low-confidence coil regions in UCSF ChimeraX^54^. The TAF1/TBP/TAF11/TAF13 subcomplex was modelled using AlphaFold2_advanced ColabFold implementation with standard settings (https://github.com/sokrypton/ColabFold/)55 and using the following protein fragments as input: TAF1 (1-300), TBP (150-339), TAF11 (50-211), TAF13 (full-length). The TAF1/TAF6^HEAT^/TAF6^HEAT^/TAF8 subcomplex was modelled using AlphaFold2 Multimer extension on COSMIC2 server with standard settings^56^ and using the following protein fragments as input: TAF1 (300-550), TAF6 (215-482), TAF8 (130-220). All structural models were visualized, analyzed and rendered in UCSF ChimeraX^54^.

## DATA AVAILABILITY

LC-MS/MS data have been deposited at PRIDE repository with the identifier PXD036358 and are publicly available as of the date of publication. This paper does not report original code. Any additional information required to reanalyze the data reported in this paper is available from the lead contact (László Tora) upon request.

## Supporting information

Supp. Tables and corresponding Refs.

## ACKNOWLEDGEMENTS

We are grateful to the IGBMC Proteomics and Photonic Microscopy platforms and cell culture facility for their assistance and instrumentation. We thank Florian Mueller for his helpful advice on imaging data. We thank Didier Devys, Stéphane Vincent and Dominique Helmlinger for thoroughly reading the manuscript and the Tora lab members for helpful discussions. This work has been supported by the *Fondation ARC pour la recherche sur le cancer* (ARCPOST-DOC2021080004113). This work was financially supported by grants from *Agence Nationale de la Recherche* (ANR) ANR-19-CE11-0003-02, ANR-PRCI-19-CE12-0029-01, ANR-20-CE12-0017-03, NIH MIRA grant (R35GM139564), and NSF (Award Number:1933344) grants. This work, as part of the ITI 2021-2028 program of the University of Strasbourg, was also supported by IdEx Unistra (ANR-10-IDEX-0002), and by SFRI-STRAT’US project (ANR 20-SFRI-0012) and EUR IMCBio (ANR-17-EURE-0023) under the framework of the French Investments for the Future Program. The research of S.A., P.K.M.S. and H.T.M.T. is financially supported by the grants from the *Deutsche Forschungsgemeinschaft* (DFG, German Research Foundation) with the project-IDs 192904750-SFB 992 and TI688/1-1.

## CONTRIBUTIONS

A.B., H.T.M.T. and L.T. conceived and designed the research. A.B., P.M., I.K., E.S., S.A., P.K.M.S. and G.Y. conducted experiments. A.B., P.M., B.M. and L.T. analyzed and interpreted the results. H.T.M.T. and L.T. supervised the study. A.B. and L.T. wrote the manuscript.

---

**Extended Data Figure 1.**
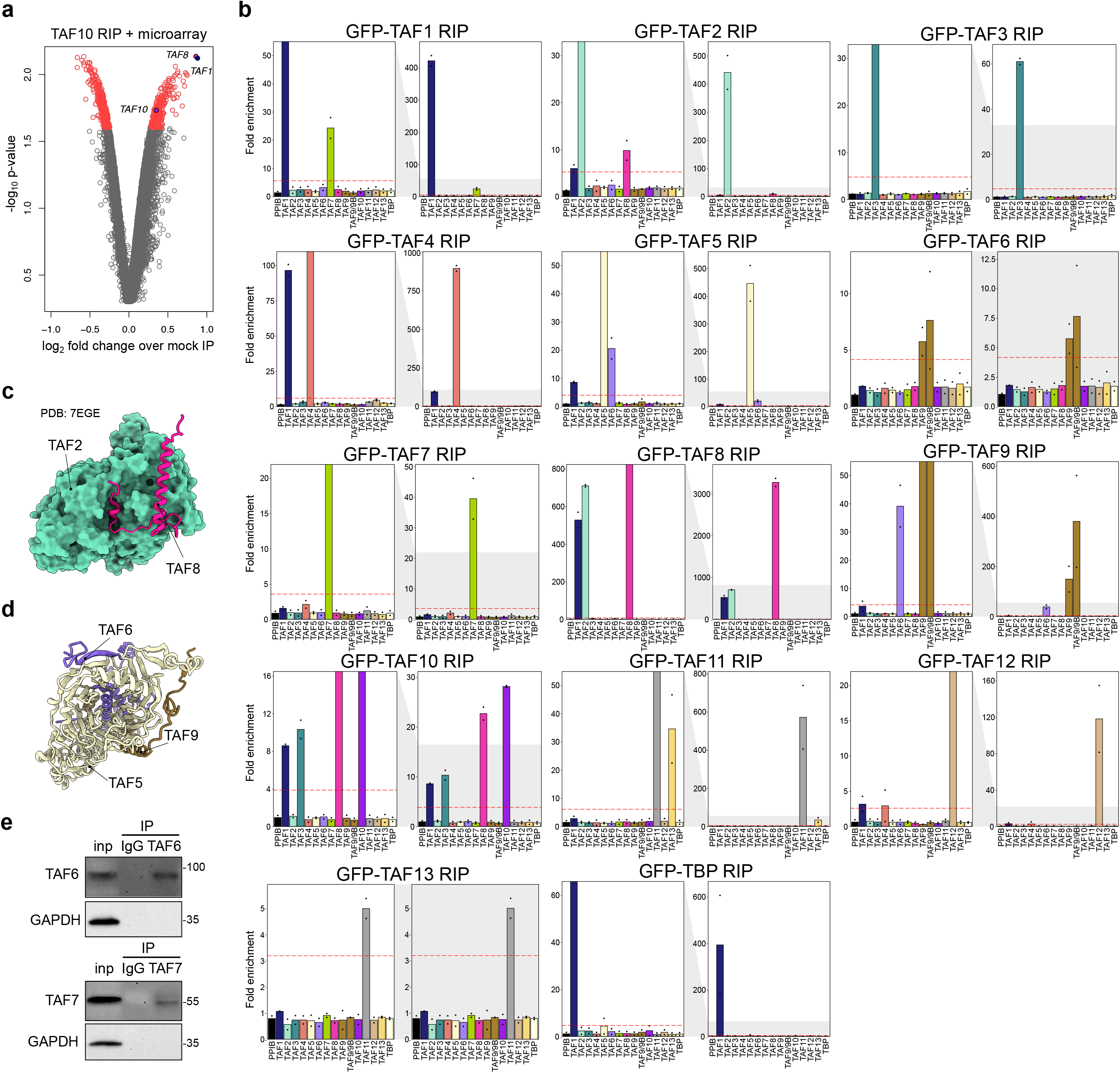
RNA immunoprecipitation (RIP) experiments to explore co-translational interactions in TFIID. **a**, Volcano plot of endogenous TAF10 RIP-microarray results. *TAF1*, *TAF8* and *TAF10* hits are highlighted. **b**, RT-qPCR results of the systematic GFP-RIP assay summarized in Fig. 1e. Each GFP-tagged TFIID subunit was used as bait in a GFP-RIP assay from polysome extracts and enrichment for TFIID subunits mRNAs was assessed by RT-qPCR. Data are expressed as mRNA fold enrichment over mock IP. For each GFP-RIP, the left panel is the zoomed version of the indicated grey-shaded area of the full-range plot (right panels). Data points correspond to biological replicas (N=2). The red dashed line threshold corresponds to 4-fold the enrichment level of the negative control target (*PPIB*). **c**, The interaction interface between TAF2/TAF8 as mapped in the TFIID Cryo-EM structure (PDB: 7EGH). The rest of TFIID subunits are not shown for clarity. **d**, Same as in (b) but for TAF5/TAF6. TAF6 completes the β-propeller blade of TAF5 WD40 domain. **e**, Western blot analysis validating the enrichment of the targeted subunit in RIP experiments against endogenous TAF6 and TAF7 from HeLa cells polysome extracts (related to Fig. 1f-g).

**Extended Data Figure 2.**
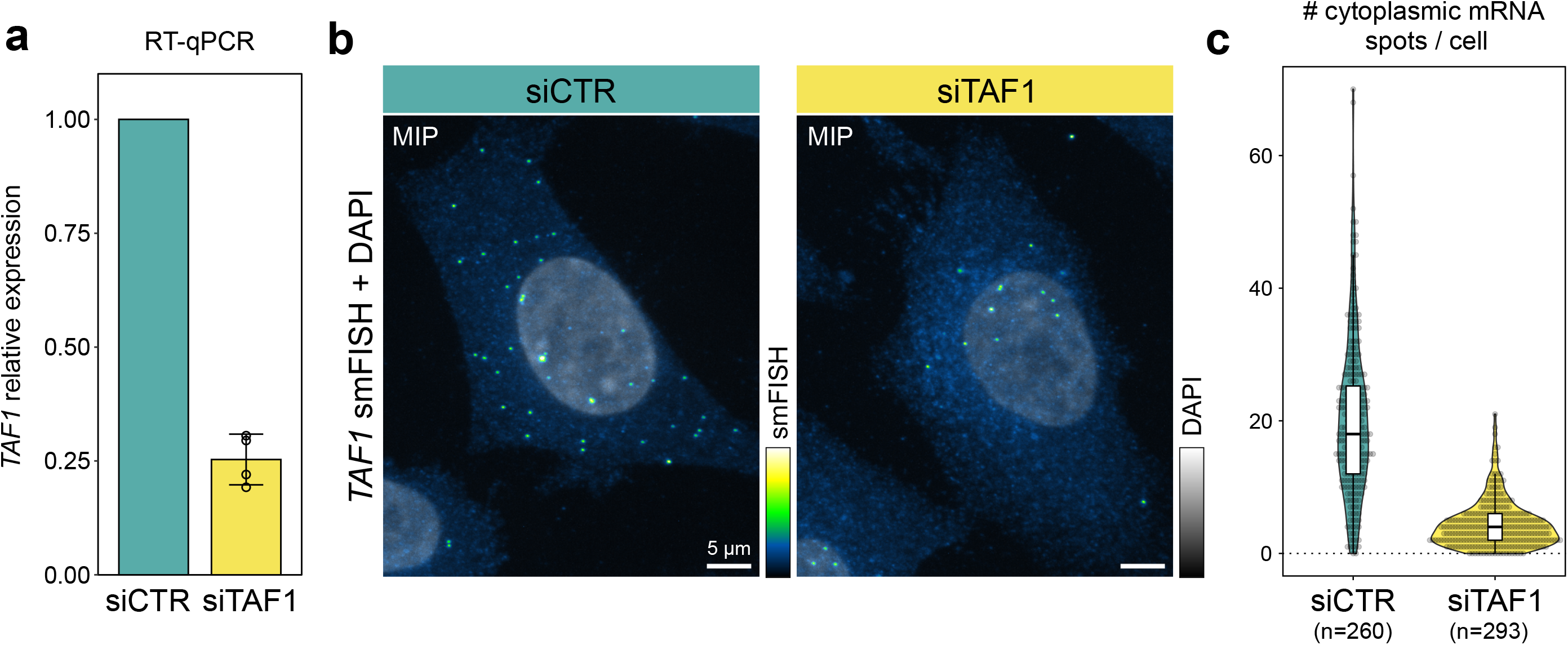
*TAF1* smFISH probes validation. **a**, *TAF1* siRNA-mediated KD assessed with RT-qPCR and expressed relative to control siRNA (CTR). Data points correspond to four biological replicas. **b**, Representative confocal maximum intensity projections (MIPs) of *TAF1* smFISH on HeLa cells transfected with control siRNA (siCTR) or siRNA directed against *TAF1* (siTAF1). smFISH and DAPI channels are displayed using the green fire blue and grayscale color scales, respectively. **c**, *TAF1* KD quantification. Violin plot representing the absolute number of cytoplasmic mRNAs per cell (total number of analyzed cells is in brackets).

**Extended Data Figure 3.**
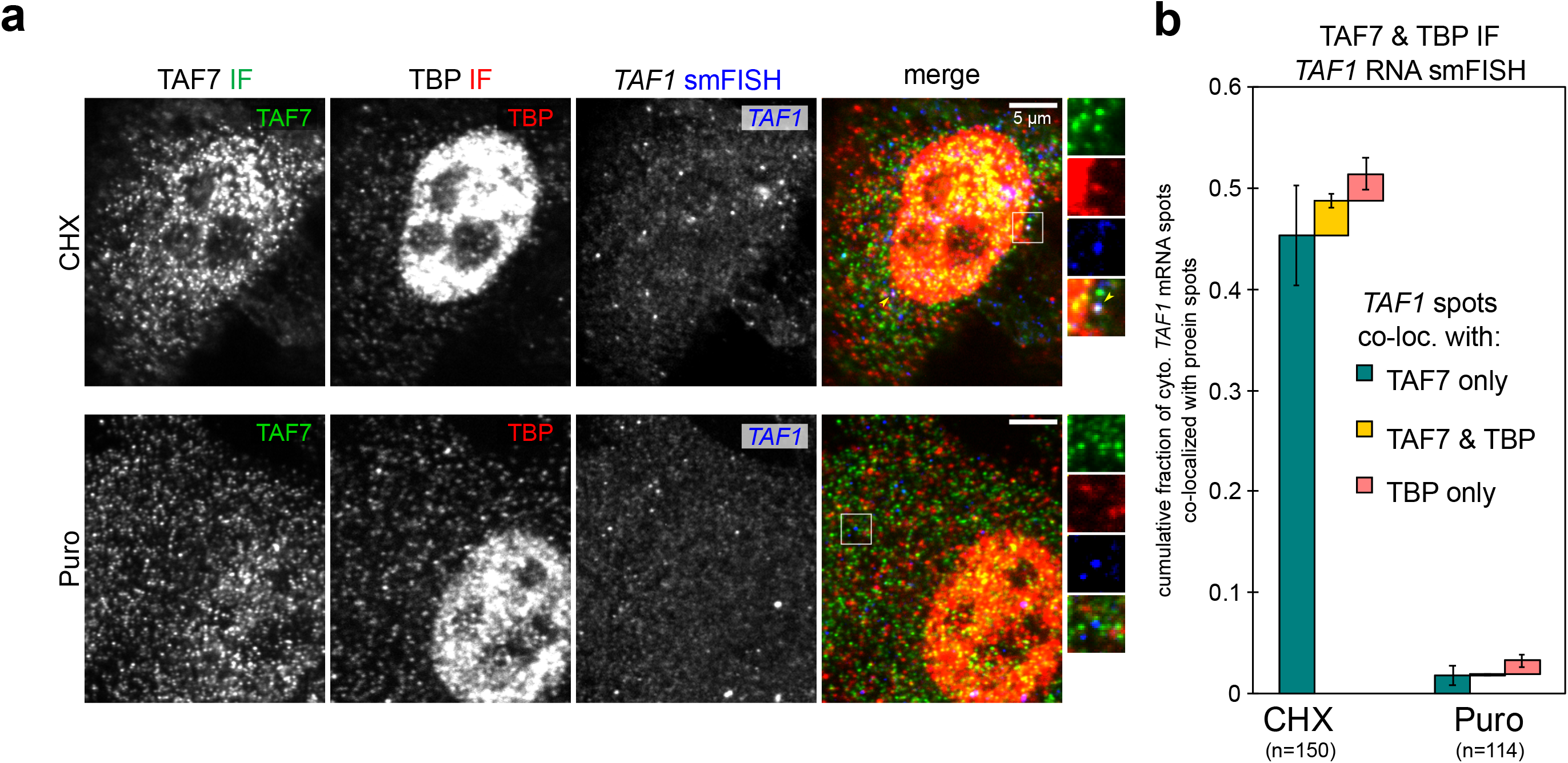
*TAF1* RNA smFISH coupled with TBP and TAF7 dual colour immunofluorescence. **a**, Representative multicolour confocal images of HeLa cells probed for TAF7 and TBP immunofluorescence (IF) coupled to *TAF1* RNA smFISH. TAF7 protein IF, TBP protein IF and *TAF1* mRNA detection in the merged image are shown in green, red and blue, respectively. Each image is a single confocal optical slice. CHX: cycloheximide; Puro: puromycin. Triple co-localized spots are indicated by yellow arrowheads. Zoom-in regions (white squares) are shown on the right. **b**, Quantification of the cumulative fraction of *TAF1* mRNAs co-localized with protein signals (TAF7, TBP or both) for each experimental condition. Bars and error bars correspond to mean and SD, respectively (N=5 for CHX; N=4 for Puro; where N corresponds to an independent field of view; total number of cells analysed is in brackets).

**Extended Data Figure 4.**
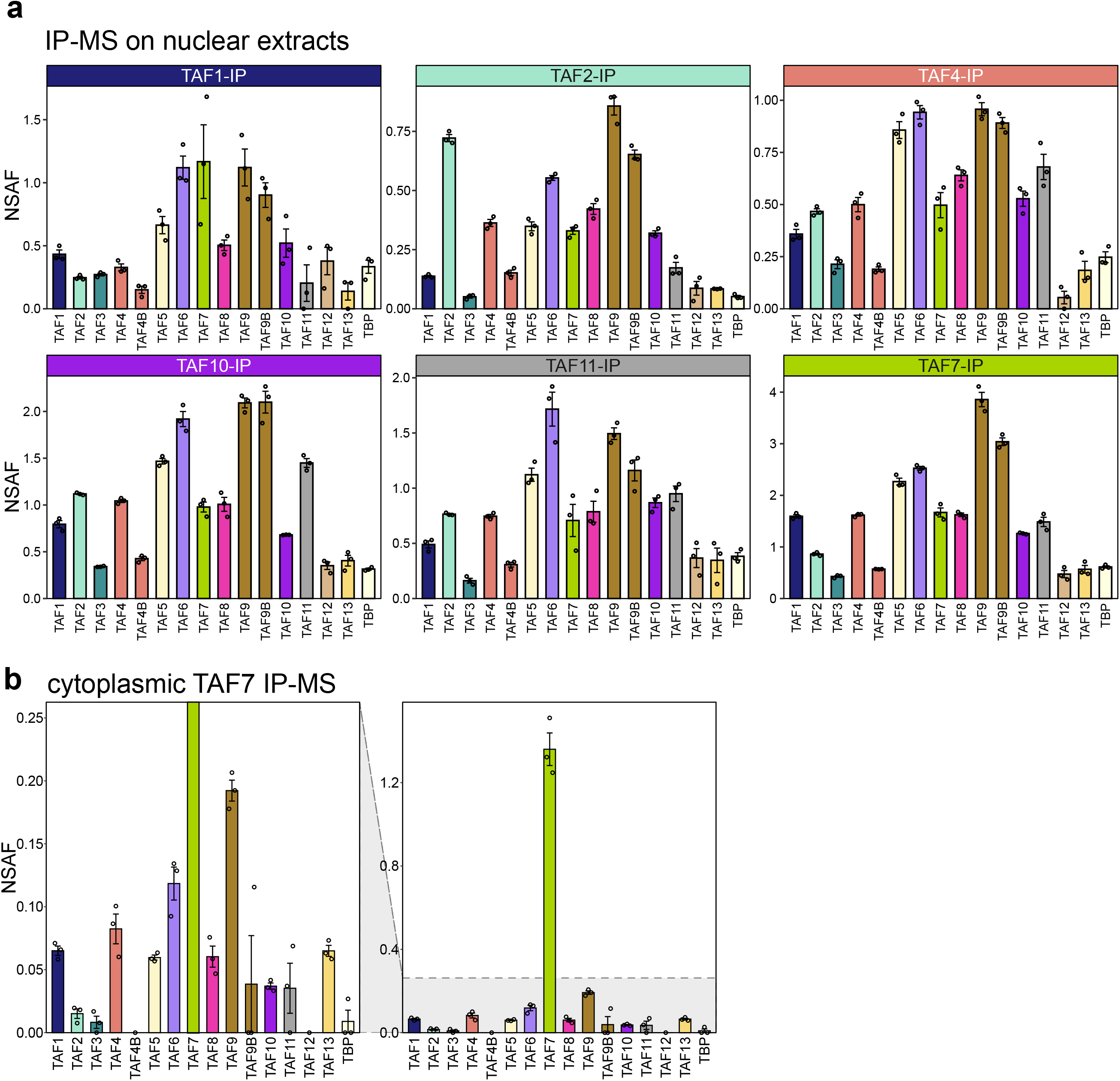
Endogenous TFIID subunits immunoprecipitation coupled to mass spectrometry. **a**, Immunoprecipitation (IP) of endogenous TFIID subunits coupled to label-free mass-spectrometry performed on human HeLa cells nuclear extracts. Bar plots represent the average NSAF (normalized spectral abundance factor) value for each detected subunit in technical triplicates. **b**, Zoomed version of cytoplasmic TAF7-IP bar plot shown in Fig. 4e to better appreciate the retrieved TFIID subunits.

**Extended Data Figure 5.**
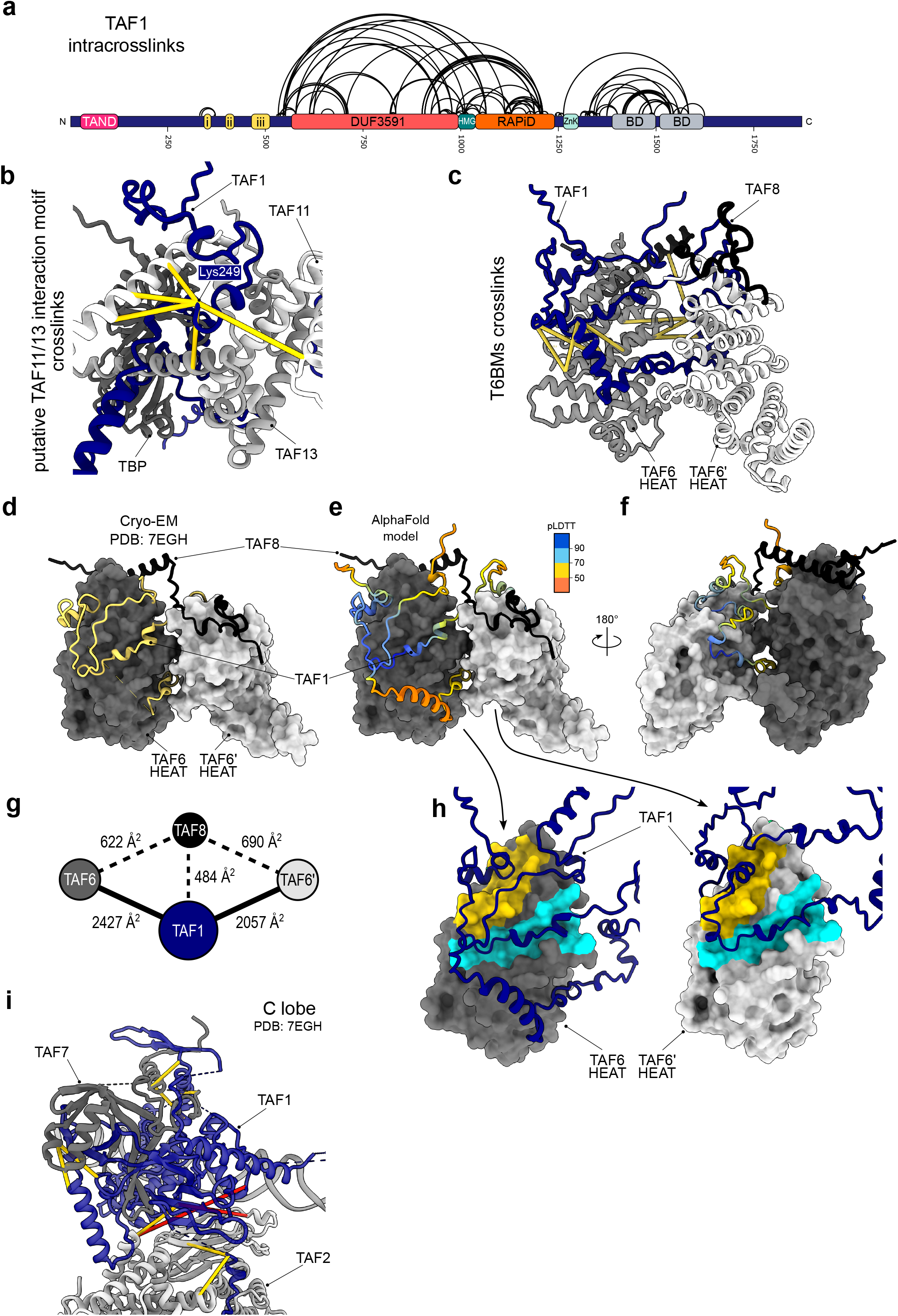
Structural insights on TAF1 interaction hotspots. **a**, Summary of TAF1 intracrosslinks from crosslinking-mass spectrometry metanalysis derived from three independent studies (Patel et al. 2018, Chen et al. 2021, Scheer et al. 2021). Only crosslinks reported in at least two independent datasets are shown. **b**, Interprotein crosslinks of TAF1 Lys249 are mapped on the AlphaFold (AF) model of the TAF1_1-300_/TAF11_50-211_/TAF13/TBP_150-339_ complex. **c**, Mapping of interprotein crosslinks between TAF1 and TAF6/TAF8 in the AF model of TAF1_300-550_/TAF6^HEAT^/TAF6^HEAT^/TAF8_128-218_ subcomplex. **d**, Cryo-EM structure of TAF1 T6BMs in complex with TAF6 HEAT domains and TAF8 (PDB: 7EGH). The rest of C lobe was removed for clarity. **e**, The AF model described in (c) is shown with TAF1 colored according to pLDDT confidence score and in the same orientation of the experimental structure shown in (d). **f**, 180 degrees rotation of the model shown in (e). **g**, Interface map of the model shown in E. The size of each node is proportional to the protein surface area. The values correspond to the buried solvent-accessible surface area between the two connected nodes. Only interfaces with a buried surface area >300 Å^2^ are shown. TAF1 bridges the two TAF6 HEAT domain copies in the complex. **h**, Equivalent surface patches contacted by TAF1 T6BMs on each of the two copies of TAF6 HEAT domains are highligthed with the same color. In each view the second HEAT domain copy is not shown for clarity. Distinct portions of TAF1 bind to equivalent surfaces on the two copies of TAF6. The representation is based on the model shown in (e). **i**, Interprotein crosslinks of TAF1 are mapped on TFIID C lobe Cryo-EM structure (PDB: 7EGH). For all panels, crosslinks compatible with crosslinker length (Cα-Cα distance < 26 Å) are displayed as yellow pseudobonds. Red pseudobonds correspond to crosslinks that exceed that distance.

**Extended Data Figure 6.**
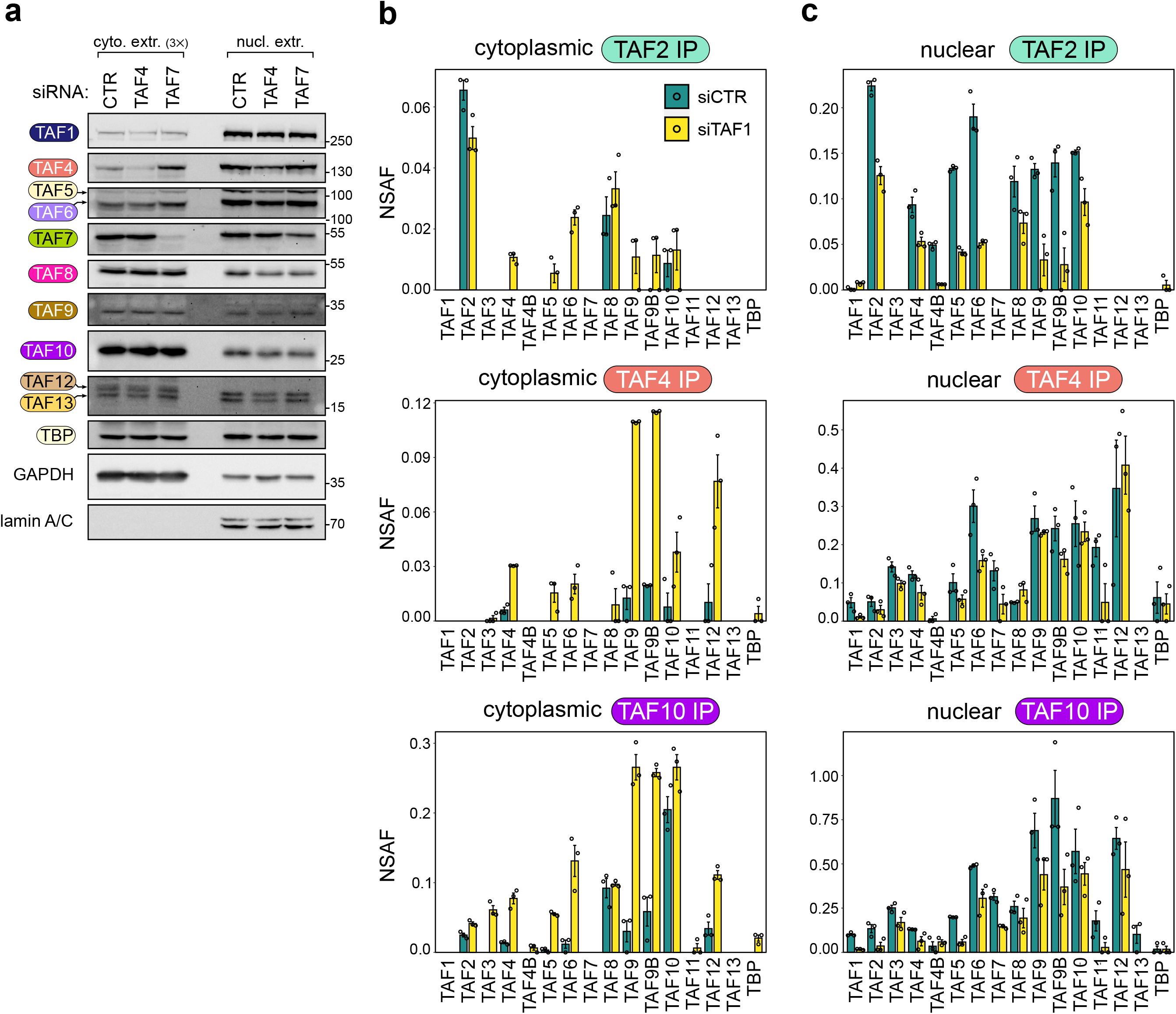
Subcellular fractionation upon TAFs knockdown. **a**, Subcellular fractionation of HeLa cells transfected with with CTR, TAF4 or TAF7 siRNAs followed by western-blot analysis of endogenous TFIID subunits distribution. GAPDH and lamin A/C were used as loading controls. The amount of loaded cytoplasmic extract is three-times the amount of the nuclear counterpart. The positions of the molecular weight markers in kDa are indicated on the right. **b**, Immunoprecipitation (IP) of endogenous TFIID subunits (TAF2, TAF4, TAF10) coupled to label-free mass-spectrometry (MS) performed on cytoplasmic extracts of HeLa cells upon *TAF1* KD. Bar plots represent the average normalized spectral abundance factor (NSAF) value for each detected subunit in technical triplicates. Error bars represent SEM. **c**, Same as in (b) but the IPs were performed on nuclear extracts. CTR: non-targeting control siRNA.

**Extended Data Figure 7.**
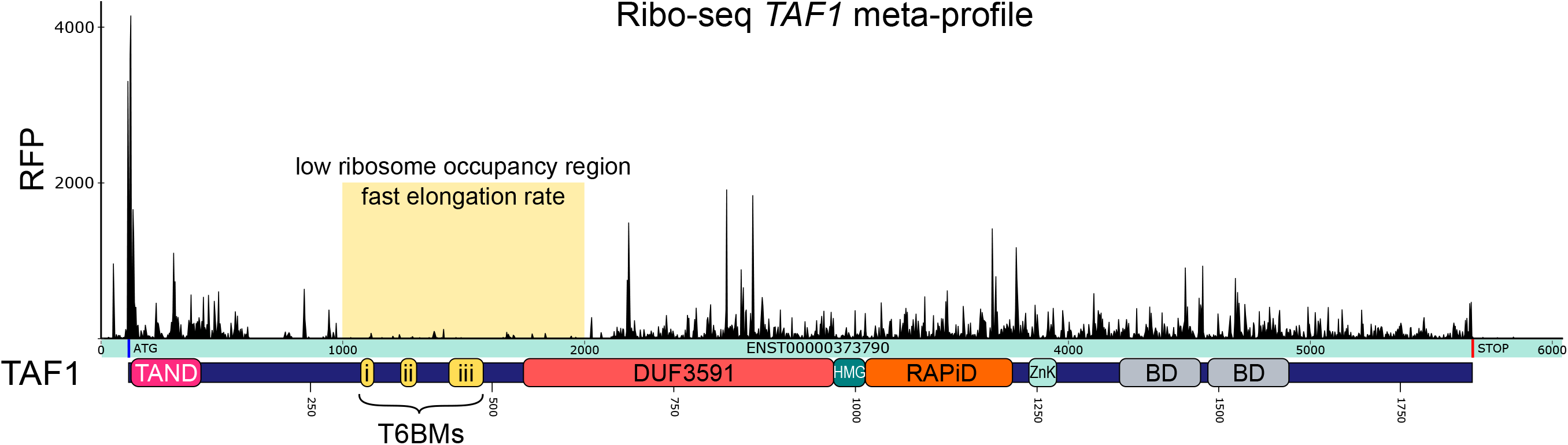
*TAF1* ribosome footprinting metaplot. Ribosome occupancy meta-profile of human *TAF1* derived from merging the available Ribo-seq datasets present in RiboCrypt browser (https://ribocrypt.org/). The yellow window highlights a region of low ribosome occupancy encompassing the three TAF6-binding motifs (T6BMs). Footprint signals coming from reading frames 2 and 3 are omitted for clarity. Below, the TAF1 functional domains are aligned to the CDS. Protein numbering matches the transcript used for this analysis (ENST00000373790). TAF1 domains are shown as in Fig. 5a. RFP, ribosome footprints.

## Notes

### Competing Interest Statement

The authors have declared no competing interest.

