## Supplementary material for "Hierarchical TAF1-dependent co-translational assembly of the basal transcription factor TFIID": Supp. Tables and corresponding Refs.

**Supplementary Table 1**: summary of TFIID co-translational assembly events. Colors correspond to subunits color-code used in this work (related to Figure 1).

| nascent protein | co-translational interaction partner |
|---|---|
| TAF1 ♦ | TAF2 ♦ |
| TAF1 ♦ | TAF4 ♦ |
| TAF1 ♦ | TAF5 ♦ |
| TAF1 ♦ | TAF6 ♦ |
| TAF1 ♦ | TAF7 ♦ |
| TAF1 ♦ | TAF8 ♦ |
| TAF1 ♦ | TAF10 ♦ |
| TAF1 ♦ | TAF12 ♦ |
| TAF1 ♦ | TBP ♦ |
| TAF2 ♦ | TAF8 ♦ |
| TAF3 ♦ | TAF10 ♦ |
| TAF4 ♦ | TAF12 ♦ |
| TAF6 ♦ | TAF9 ♦ |
| TAF6 ♦ | TAF5 ♦ |
| TAF7 ♦ | TAF1 ♦ |
| TAF8 ♦ | TAF2 ♦ |
| TAF8 ♦ | TAF10 ♦ |
| TAF9 ♦ | TAF6 ♦ |
| TAF11 ♦ | TAF13 ♦ |
| TAF13 ♦ | TAF11 ♦ |

**Supplementary Table 2**: X-linking MS combined table (related to Figure 5)

Summary of TAF1-centered crosslinking-MS metanalysis performed on seven distinct datasets from three different groups: Patel et al., 2018 (TFIID), Scheer et al., 2021 (TFIID), Chen et al., 2021 (TFIID incorporated in different preinitiation complexes: cPICscp, cPICpuma, mPICscp, hPICscp, p53hPIChdm2). Only crosslinks found in more than one dataset were considered.

**TAF1 interprotein crosslinks**

| Protein 1 | Protein 2 | Position 1 | Position 2 | Protein 1 | Protein 2 | Position 1 | Position 2 |
|---|---|---|---|---|---|---|---|
| TAF1 | TAF11 | 249 | 95 | TAF1 | TAF6 | 378 | 357 |
| TAF1 | TAF11 | 249 | 97 | TAF1 | TAF6 | 424 | 342 |
| TAF1 | TAF11 | 249 | 105 | TAF1 | TAF6 | 427 | 342 |
| TAF1 | TAF11 | 249 | 197 | TAF1 | TAF6 | 427 | 287 |
| TAF1 | TAF11 | 249 | 197 | TAF1 | TAF7 | 622 | 5 |
| TAF1 | TAF13 | 249 | 101 | TAF1 | TAF7 | 622 | 291 |
| TAF1 | TAF13 | 249 | 101 | TAF1 | TAF7 | 674 | 40 |
| TAF1 | TAF2 | 544 | 383 | TAF1 | TAF7 | 674 | 40 |
| TAF1 | TAF2 | 544 | 788 | TAF1 | TAF7 | 817 | 5 |
| TAF1 | TAF2 | 562 | 788 | TAF1 | TAF7 | 819 | 5 |
| TAF1 | TAF2 | 576 | 384 | TAF1 | TAF7 | 832 | 5 |
| TAF1 | TAF2 | 576 | 565 | TAF1 | TAF7 | 889 | 164 |
| TAF1 | TAF2 | 701 | 513 | TAF1 | TAF7 | 906 | 164 |
| TAF1 | TAF2 | 710 | 595 | TAF1 | TAF7 | 906 | 167 |
| TAF1 | TAF2 | 945 | 513 | TAF1 | TAF7 | 1127 | 164 |
| TAF1 | TAF5 | 370 | 318 | TAF1 | TAF7 | 1187 | 164 |
| TAF1 | TAF6 | 330 | 65 | TAF1 | TAF7 | 1201 | 155 |
| TAF1 | TAF6 | 330 | 196 | TAF1 | TAF7 | 1201 | 153 |
| TAF1 | TAF6 | 330 | 158 | TAF1 | TAF7 | 1208 | 167 |
| TAF1 | TAF6 | 335 | 196 | TAF1 | TAF8 | 427 | 178 |
| TAF1 | TAF6 | 370 | 196 | TAF1 | TAF9 | 330 | 24 |
| TAF1 | TAF6 | 370 | 367 | TAF1 | TAF9 | 330 | 10 |
| TAF1 | TAF6 | 370 | 361 | TAF1 | TBP | 168 | 243 |
| TAF1 | TAF6 | 378 | 361 | TAF1 | TBP | 170 | 333 |
| TAF1 | TAF6 | 378 | 361 | TAF1 | TBP | 1009 | 181 |
| TAF1 | TAF6 | 378 | 367 | | | | |

**TAF1 intraprotein crosslinks**

| Protein 1 | Protein 2 | Position 1 | Position 2 | Protein 1 | Protein 2 | Position 1 | Position 2 |
|---|---|---|---|---|---|---|---|
| TAF1 | TAF1 | 330 | 370 | TAF1 | TAF1 | 1009 | 1046 |
| TAF1 | TAF1 | 335 | 370 | TAF1 | TAF1 | 1009 | 1127 |
| TAF1 | TAF1 | 367 | 370 | TAF1 | TAF1 | 1009 | 1111 |
| TAF1 | TAF1 | 527 | 531 | TAF1 | TAF1 | 1046 | 1063 |
| TAF1 | TAF1 | 531 | 536 | TAF1 | TAF1 | 1046 | 1201 |
| TAF1 | TAF1 | 531 | 1436 | TAF1 | TAF1 | 1111 | 1127 |
| TAF1 | TAF1 | 531 | 976 | TAF1 | TAF1 | 1111 | 1201 |
| TAF1 | TAF1 | 531 | 544 | TAF1 | TAF1 | 1111 | 1187 |
| TAF1 | TAF1 | 536 | 544 | TAF1 | TAF1 | 1112 | 1187 |
| TAF1 | TAF1 | 536 | 549 | TAF1 | TAF1 | 1112 | 1201 |
| TAF1 | TAF1 | 536 | 576 | TAF1 | TAF1 | 1112 | 1127 |
| TAF1 | TAF1 | 544 | 943 | TAF1 | TAF1 | 1112 | 1177 |

| | | | | | | | |
|---|---|---|---|---|---|---|---|
| TAF1 | TAF1 | 544 | 1063 | TAF1 | TAF1 | 1112 | 1208 |
| TAF1 | TAF1 | 544 | 1009 | TAF1 | TAF1 | 1117 | 1127 |
| TAF1 | TAF1 | 544 | 576 | TAF1 | TAF1 | 1127 | 1208 |
| TAF1 | TAF1 | 544 | 549 | TAF1 | TAF1 | 1127 | 1187 |
| TAF1 | TAF1 | 544 | 710 | TAF1 | TAF1 | 1127 | 1166 |
| TAF1 | TAF1 | 549 | 706 | TAF1 | TAF1 | 1127 | 1201 |
| TAF1 | TAF1 | 549 | 707 | TAF1 | TAF1 | 1166 | 1187 |
| TAF1 | TAF1 | 549 | 1063 | TAF1 | TAF1 | 1187 | 1208 |
| TAF1 | TAF1 | 549 | 705 | TAF1 | TAF1 | 1201 | 1208 |
| TAF1 | TAF1 | 576 | 701 | TAF1 | TAF1 | 1201 | 1222 |
| TAF1 | TAF1 | 576 | 1063 | TAF1 | TAF1 | 1240 | 1255 |
| TAF1 | TAF1 | 576 | 943 | TAF1 | TAF1 | 1240 | 1244 |
| TAF1 | TAF1 | 576 | 945 | TAF1 | TAF1 | 1249 | 1255 |
| TAF1 | TAF1 | 611 | 832 | TAF1 | TAF1 | 1249 | 1261 |
| TAF1 | TAF1 | 611 | 621 | TAF1 | TAF1 | 1261 | 1581 |
| TAF1 | TAF1 | 611 | 622 | TAF1 | TAF1 | 1305 | 1322 |
| TAF1 | TAF1 | 611 | 976 | TAF1 | TAF1 | 1305 | 1327 |
| TAF1 | TAF1 | 621 | 832 | TAF1 | TAF1 | 1317 | 1339 |
| TAF1 | TAF1 | 622 | 1111 | TAF1 | TAF1 | 1317 | 1327 |
| TAF1 | TAF1 | 641 | 674 | TAF1 | TAF1 | 1322 | 1327 |
| TAF1 | TAF1 | 701 | 710 | TAF1 | TAF1 | 1322 | 1344 |
| TAF1 | TAF1 | 701 | 707 | TAF1 | TAF1 | 1322 | 1339 |
| TAF1 | TAF1 | 705 | 1063 | TAF1 | TAF1 | 1327 | 1344 |
| TAF1 | TAF1 | 705 | 710 | TAF1 | TAF1 | 1327 | 1535 |
| TAF1 | TAF1 | 705 | 707 | TAF1 | TAF1 | 1327 | 1487 |
| TAF1 | TAF1 | 817 | 832 | TAF1 | TAF1 | 1327 | 1329 |
| TAF1 | TAF1 | 817 | 1004 | TAF1 | TAF1 | 1339 | 1581 |
| TAF1 | TAF1 | 819 | 832 | TAF1 | TAF1 | 1339 | 1347 |
| TAF1 | TAF1 | 819 | 1004 | TAF1 | TAF1 | 1344 | 1487 |
| TAF1 | TAF1 | 831 | 1205 | TAF1 | TAF1 | 1344 | 1354 |
| TAF1 | TAF1 | 899 | 1208 | TAF1 | TAF1 | 1344 | 1399 |
| TAF1 | TAF1 | 899 | 1187 | TAF1 | TAF1 | 1353 | 1399 |
| TAF1 | TAF1 | 899 | 1201 | TAF1 | TAF1 | 1372 | 1487 |
| TAF1 | TAF1 | 906 | 1208 | TAF1 | TAF1 | 1372 | 1555 |
| TAF1 | TAF1 | 906 | 1201 | TAF1 | TAF1 | 1412 | 1419 |
| TAF1 | TAF1 | 906 | 1187 | TAF1 | TAF1 | 1412 | 1535 |
| TAF1 | TAF1 | 922 | 1009 | TAF1 | TAF1 | 1412 | 1542 |
| TAF1 | TAF1 | 922 | 1201 | TAF1 | TAF1 | 1414 | 1419 |
| TAF1 | TAF1 | 943 | 971 | TAF1 | TAF1 | 1415 | 1535 |
| TAF1 | TAF1 | 943 | 967 | TAF1 | TAF1 | 1415 | 1542 |
| TAF1 | TAF1 | 945 | 976 | TAF1 | TAF1 | 1419 | 1534 |
| TAF1 | TAF1 | 967 | 976 | TAF1 | TAF1 | 1419 | 1542 |
| TAF1 | TAF1 | 971 | 979 | TAF1 | TAF1 | 1419 | 1535 |
| TAF1 | TAF1 | 971 | 986 | TAF1 | TAF1 | 1454 | 1534 |
| TAF1 | TAF1 | 976 | 979 | TAF1 | TAF1 | 1454 | 1535 |
| TAF1 | TAF1 | 976 | 986 | TAF1 | TAF1 | 1463 | 1535 |
| TAF1 | TAF1 | 979 | 987 | TAF1 | TAF1 | 1463 | 1534 |

| | | | | | | | |
|---|---|---|---|---|---|---|---|
| TAF1 | TAF1 | 979 | 986 | TAF1 | TAF1 | 1480 | 1493 |
| TAF1 | TAF1 | 986 | 1009 | TAF1 | TAF1 | 1480 | 1487 |
| TAF1 | TAF1 | 986 | 1046 | TAF1 | TAF1 | 1482 | 1487 |
| TAF1 | TAF1 | 986 | 1127 | TAF1 | TAF1 | 1482 | 1493 |
| TAF1 | TAF1 | 986 | 1001 | TAF1 | TAF1 | 1487 | 1493 |
| TAF1 | TAF1 | 1001 | 1009 | TAF1 | TAF1 | 1493 | 1559 |
| TAF1 | TAF1 | 1001 | 1018 | TAF1 | TAF1 | 1534 | 1542 |
| TAF1 | TAF1 | 1004 | 1018 | TAF1 | TAF1 | 1542 | 1581 |
| TAF1 | TAF1 | 1009 | 1063 | TAF1 | TAF1 | 1561 | 1622 |
| TAF1 | TAF1 | 1009 | 1006 | | | | |

**Supplementary Table 3**: oligonucleotide sequences and antibodies

| | |
|---|---|
| **RT-qPCR primers** | |
| GAPDH Fwd | TCGACAGTCAGCCGCATCTTCTTT |
| GAPDH Rev | ACCAAATCCGTTGACTCCGACCTT |
| PPIB Fwd | CCGAACGCAACATGAAGGTG |
| PPIB Rev | ACCAAAGATCACCCGGCCTA |
| TAF1 Fwd | TTCCAACCCTGTTGCCATGA |
| TAF1 Rev | TTTCTGCGAACCTCATCCGC |
| TAF2 Fwd | CATGTGTACCGCCAAAGT |
| TAF2 Rev | GCAGTTGCTTCTGTGTAAATC |
| TAF3 Fwd | GACGACTGCGATGACTGGTA |
| TAF3 Rev | CTTCTTGTTCGCACACTTGG |
| TAF4 Fwd | GCCGCGCAAACTTTGAATG |
| TAF4 Rev | TTGTTGACCAGGCTGACAGC |
| TAF5 Fwd | AGTTGGAAGTGTTGCTGTGG |
| TAF5 Rev | TCCTTGTTGGTTGTAGGCTGAC |
| TAF6 Fwd | CCAGGAGTTCATTCCTTTCC |
| TAF6 Rev | TGATGTCGCTCAGATCAACC |
| TAF7 Fwd | TCTACTGTGAGAAGGGCAGTAC |
| TAF7 Rev | ATTCCATGACGCCCATCAGG |
| TAF8 Fwd | ACAGAGGCAGGGTTTGAGAGT |
| TAF8 Rev | AGACTTGGCACTTCTCCCAAT |
| TAF9 Fwd | GGAGTTTGCCTTCCGATATG |
| TAF9 Rev | CGCACATCATCTGCATCAAC |
| TAF9/9B Fwd | ATCAAACCCCTTTGCCA |
| TAF9/9B Rev | TTCAGCCTATAGTTTGGAGC |
| TAF10 Fwd | TGCCAATGATGCCCTACAGC |
| TAF10 Rev | AGGGCAGGGGTCAAGTCCTC |
| TAF11 Fwd | AAAGGCTGATCCAGTCCATCAC |
| TAF11 Rev | TTTCTCCCCACTTCTCACACAC |
| TAF12 Fwd | TATGAGGACCCGCACTCCTAC |
| TAF12 Rev | GCCGAGCTTTGGACTTCAGC |
| TAF13 Fwd | AATTGGAGGAGGTGCAGAAGG |
| TAF13 Rev | TGGTCATCCCCAAAGCCATAC |
| TBP Fwd | TCATACCGTGCTGCTATCT |
| TBP Rev | CTCCCTCAAACCAACTTGTC |
| Rplp0-mouse Fwd | TTCTGAGTGATGTGCAGCTG |
| Rplp0-mouse Rev | GGAGATGTTCAGCATGTTCAGC |
| Taf1-mouse Fwd | TGGAGATGGTGATCTTGCAG |
| Taf1-mouse Rev | TCCTCATCATCTTCGCCTTC |
| Taf8-mouse Fwd | ATATCAGCACGGACGATTCC |
| Taf8-mouse Rev | GGTTATCGATGACGCTCTCC |
| Taf10-mouse Fwd | CCACGCATAATTCGGCTCAT |
| Taf10-mouse Rev | CCTCCATGGTTAGGTGTACT |
| **smiFISH primary probes** (including FLAP extension) | |
| CTNNB1_01 | CTCATGTTCCATCATGGGGTCCATACCTTACACTCGGACCTCGTCGACATGCATT |
| CTNNB1_02 | GCATCCTGGCCATATCCACCAGAGTGTTACACTCGGACCTCGTCGACATGCATT |
| CTNNB1_03 | TGTTCTGAAGAGAGAGCTGGTCAGCTCAACTTTACACTCGGACCTCGTCGACATGCATT |
| CTNNB1_04 | GCCGTTTCTTGTAATCTTGTGGCTTGTCCTTTACACTCGGACCTCGTCGACATGCATT |
| CTNNB1_05 | AGCTGTGGCTCCCTCAGCTTCAATAGTTACACTCGGACCTCGTCGACATGCATT |

| | |
|---|---|
| CTNNB1_06 | TGCAGCTTCCTTGTCCTGAGCAAGTTCATTACACTCGGACCTCGTCGACATGCATT |
| CTNNB1_07 | GAGCTAGGATGTGAAGGGCTCCGGTACAACTTACACTCGGACCTCGTCGACATGCATT |
| CTNNB1_08 | AAATTGCTGCTGTGTCCCACCCATGGTTACACTCGGACCTCGTCGACATGCATT |
| CTNNB1_09 | GGCCAGTGGGATGGTGGGTGTAAGAGCTTACACTCGGACCTCGTCGACATGCATT |
| CTNNB1_10 | TGGGCCATCTCTGCTTCTTGGTGTCGTTACACTCGGACCTCGTCGACATGCATT |
| CTNNB1_11 | TGATGTCTTCCCTGTCACCAGCCCGATTACACTCGGACCTCGTCGACATGCATT |
| CTNNB1_12 | GTCCCAAGGAGACCTTCCATCCCTTCTTACACTCGGACCTCGTCGACATGCATT |
| CTNNB1_13 | AGCACCTTCAGCACTCTGCTTGTGGTTTACACTCGGACCTCGTCGACATGCATT |
| CTNNB1_14 | ACCACTAGCCAGTATGATGAGCTTGCTTTTACACTCGGACCTCGTCGACATGCATT |
| CTNNB1_15 | TTGTTTTGTTGAGCAAGGCAACCATTTTCTGCTTACACTCGGACCTCGTCGACATGCATT |
| CTNNB1_16 | TGGGAAAGGTTATGCAAGGTCCCAGCGGTATTACACTCGGACCTCGTCGACATGCATT |
| CTNNB1_17 | ATAGCGTGTCTGGAAGCTTCCTTTTTAGAAAGTTACACTCGGACCTCGTCGACATGCATT |
| CTNNB1_18 | TGGTCCTCGTCATTTAGCAGTTTTGTCAGTTCTTACACTCGGACCTCGTCGACATGCATT |
| CTNNB1_19 | ATTGCACGTGTGGCAAGTTCTGCATCATCTTACACTCGGACCTCGTCGACATGCATT |
| CTNNB1_20 | ATGGTTCAGCCAAACGCTGGACATTAGTGGTTACACTCGGACCTCGTCGACATGCATT |
| CTNNB1_21 | GTCCATCAATATCAGCTACTTGTTCTTGAGTGTTACACTCGGACCTCGTCGACATGCATT |
| CTNNB1_22 | CTTGGGAGGTATCCACATCCTCTTCCTTTACACTCGGACCTCGTCGACATGCATT |
| CTNNB1_23 | ATTGCCTTTACCACTCAGAGAAGGAGCTGTTTACACTCGGACCTCGTCGACATGCATT |
| CTNNB1_24 | GTGGCACCAGAATGGATTCCAGAGTCCAGTTACACTCGGACCTCGTCGACATGCATT |
| TAF1_01 | GCAATGGAGTGGAAATCCTCACTGTCCTTTACACTCGGACCTCGTCGACATGCATT |
| TAF1_02 | CTCATAGCTCCCATAACTGATGTTGCTATTTACACTCGGACCTCGTCGACATGCATT |
| TAF1_03 | TCCACTTTCACTCAGCTGGATAGCAGAGTTACACTCGGACCTCGTCGACATGCATT |
| TAF1_04 | GGAGATGTTCTTACGTATGGTCTCTAAATCCATTACACTCGGACCTCGTCGACATGCATT |
| TAF1_05 | AGCAAGGGGTTGATAGCTTTCTCTAAGCGAGTTACACTCGGACCTCGTCGACATGCATT |
| TAF1_06 | AGTCCTTTACAACCTTTGCATTGACTGGAGTTTACACTCGGACCTCGTCGACATGCATT |
| TAF1_07 | AAGGTGGCGCATTTGTTTGATAATAGAGGGGGTTACACTCGGACCTCGTCGACATGCATT |
| TAF1_08 | GAATTTGTTAGTCCTCATGTGTCCAATGGCATTACACTCGGACCTCGTCGACATGCATT |
| TAF1_09 | ACATAGGCATCAATGACAGCTGGTTTTCGGTTACACTCGGACCTCGTCGACATGCATT |
| TAF1_10 | AGACTTCAGTTGATGACAGAACCTTGTTCTGTTTACACTCGGACCTCGTCGACATGCATT |
| TAF1_11 | GTCAAAGATGCGCTGACATTCCTCTTTGTAATTACACTCGGACCTCGTCGACATGCATT |
| TAF1_12 | AACTTTTTAATCTCTTCCTCAGGCACACCTTACACTCGGACCTCGTCGACATGCATT |
| TAF1_13 | AGAAGTTGCTTGGCATTTTTCAGGGAAAGGCGTTACACTCGGACCTCGTCGACATGCATT |
| TAF1_14 | AGCAATGAAGGCCCTTGTGGTGTTCCAAGTTACACTCGGACCTCGTCGACATGCATT |
| TAF1_15 | GCAGTGCGAACTTCATCATCAATCTTCATTTACACTCGGACCTCGTCGACATGCATT |
| TAF1_16 | AATTCCCGAATATAGTAACCCTGTCTTGTCCTTACACTCGGACCTCGTCGACATGCATT |
| TAF1_17 | ATGTGGGTGGGAAAGAAGGGCTGCCGTAATTTACACTCGGACCTCGTCGACATGCATT |
| TAF1_18 | GGCATGGCCTGAGCATCCCAAATGATTTACACTCGGACCTCGTCGACATGCATT |
| TAF1_19 | CCAGCGTCCATATACCAGATCCTCATTTACACTCGGACCTCGTCGACATGCATT |
| TAF1_20 | GATGATATCATCCTCCCAATGCAGCTTTACACTCGGACCTCGTCGACATGCATT |
| TAF1_21 | TCGTGATTTCATCATCAGAGAGACACTGCTTTACACTCGGACCTCGTCGACATGCATT |
| TAF1_22 | CAAAGACTTCTGGCTGACTTCTGATTCTACTTACACTCGGACCTCGTCGACATGCATT |
| TAF1_23 | CAATGGAAGGGTCAGCTTTCCATCTTCAGATTACACTCGGACCTCGTCGACATGCATT |

| | | | |
|---|---|---|---|
| TAF1_24 | TTCATCATTTACCAAGGCACCGTCAGTCCTTACACTCGGACCTCGTCGACATGCATT | | |
| **smiFISH secondary FLAP probe** | | | |
| 2×Cy3-FLAP | AATGCATGTCGACGAGGTCCGAGTGTAA | | |
| **Antibodies** | | | |
| Target | Clonality | Reference/Clone | Application |
| GST | mouse mAb | 15TF2 1D10 (Creative Biolabs) | IP |
| TAF1 | rabbit pAb | ab188427 (Abcam) | IF |
| TAF1 | rabbit pAb | ab264327 (Abcam) | IP, WB |
| TAF2 | rabbit pAb | #3038 (Trowitzsch et al., 2015) | IP |
| TAF4 | mouse mAb | 32TA 2B9 (Mohan et al., 2003) | IP, IF, WB |
| TAF5 | mouse mAb | 1TA 1C2 (Dantonel et al., 1997) | WB |
| TAF6 | mouse mAb | 25TA 2G7 (Dantonel et al., 1997) | WB |
| TAF6 | rabbit pAb | A301-275A (Bethyl) | RIP |
| TAF7 | rabbit pAb | #3475 (Bardot et al., 2017) | IP, IF |
| TAF7 | mouse mAb | 31TA 2C12 (present work) | RIP |
| TAF7 | mouse mAb | 19TA 2C7 (Lavigne et al., 1996) | WB |
| TAF8 | rabbit pAb | #3478 (Bardot et al., 2017) | WB |
| TAF9 | goat pAb | sc-1248 (Santa Cruz Biotechnology) | WB |
| TAF10 | mouse mAb | 23TA 1H8 (Soutoglou et al., 2005) | IP, RIP |
| TAF10 | mouse mAb | 6TA 2B11 (Wieczorek et al., 1998) | RIP, IF, WB |
| TAF11 | mouse mAb | 15TA 2B4 (Gupta et al., 2017) | IP |
| TAF12 | mouse mAb | 22TA 2A1 (Brand et al., 2001) | WB |
| TAF13 | mouse mAb | 16TA 3C12 (Mengus et al., 1995) | WB |
| TBP | mouse mAb | 3TF1 3G3 (Brou et al., 1993) | WB, IF |
| SUPT7L | rabbit pAb | A302-803A (Bethyl) | IF |
| lamin A/C | mouse mAb | sc-7292 (Santa Cruz Biotechnology) | WB |
| GAPDH | rabbit mAb | 14C10 (Cell Signaling Technology) | WB |
| histone H3 | rabbit pAb | ab1791 (Abcam) | WB |
